# Identification of evolutionary trajectories shared across human betacoronaviruses

**DOI:** 10.1101/2021.05.24.445313

**Authors:** Marina Escalera-Zamudio, Sergei L. Kosakovsky Pond, Natalia Martínez de la Viña, Bernardo Gutiérrez, Rhys P. D. Inward, Julien Thézé, Lucy van Dorp, Hugo G. Castelán-Sánchez, Thomas A. Bowden, Oliver G. Pybus, Ruben J.G. Hulswit

**Author notes:** Email addresses.

## Abstract

Comparing the evolution of distantly related viruses can provide insights into common adaptive processes related to shared ecological niches. Phylogenetic approaches, coupled with other molecular evolution tools, can help identify mutations informative on adaptation, whilst the structural contextualization of these to functional sites of proteins may help gain insight into their biological properties. Two zoonotic betacoronaviruses capable of sustained human-to-human transmission have caused pandemics in recent times (SARS-CoV-1 and SARS-CoV-2), whilst a third virus (MERS-CoV) is responsible for sporadic outbreaks linked to animal infections. Moreover, two other betacoronaviruses have circulated endemically in humans for decades (HKU1 and OC43). To search for evidence of adaptive convergence between established and emerging betacoronaviruses capable of sustained human-to-human transmission (HKU1, OC43, SARS-CoV-1 and SARS-CoV-2), we developed a methodological pipeline to classify shared non-synonymous mutations as putatively denoting homoplasy (repeated mutations that do not share direct common ancestry) or stepwise evolution (sequential mutations leading towards a novel genotype). In parallel, we look for evidence of positive selection, and draw upon protein structure data to identify potential biological implications. We find 30 mutations, with four of these [codon sites 18121 (nsp14/residue 28), 21623 (spike/21), 21635 (spike/25) and 23948 (spike/796); SARS-CoV-2 genome numbering] displaying evolution under positive selection and proximity to functional protein regions. Our findings shed light on potential mechanisms underlying betacoronavirus adaptation to the human host and pinpoint common mutational pathways that may occur during establishment of human endemicity.

## INTRODUCTION

Mutation is a fundamental process of evolution, and a main source of variability enabling genetic diversity (Loewe and Hill 2010). However, most mutations are expected to be either eliminated through purifying selection, or to be selectively neutral and thus fixed randomly. Although RNA viruses change rapidly due to their relatively small genomes and high mutation rates, the mutational pathways leading to adaptation are often limited by functional and evolutionary constraints (such as epistasis, referring to the adaptive effect of a mutation being dependent on the genetic background in which it appears) (Dolan et al. 2018). Following a nearly-neutral evolutionary theory, only the smallest proportion of mutations are expected to be adaptive and subsequently fixed through positive selection (Ohta 1973, Pond et al. 2012). Thus, given a reduced admissible genetic variability in viruses, evolutionary trajectories are limited, and may sometimes display recurrent mutational patterns denoting adaptive convergence, at least for viruses sharing ecological niches (Gutierrez, Escalera-Zamudio, and Oliver G Pybus 2019).

Coronaviruses are well known for their propensity to switch host species, evidenced by multiple zoonotic introductions into the human population. There are four betacoronavirus species capable of sustained human transmission: OC43, HKU1, SARS-CoV-1, and SARS-CoV-2. These viruses are further classified into two subgenera: *Embecovirus* and *Sarbecovirus*. The *Embecovirus* subgenus includes OC43 first described in 1967 (McIntosh et al. 1967), and HKU1 identified in 2005 (Woo et al. 2005). Both were introduced into the human population through independent zoonotic events estimated to have occurred at least 50 years ago (Vijgen et al. 2005; Lau et al. 2015; Kistler and Bedford 2021; Pollett et al. 2021), and are associated with mild respiratory disease (Su et al. 2016). On the other hand, the two viruses belonging to the *Sarbecovirus* subgenus (SARS-CoV-1 and SARS-CoV-2) were introduced independently but much more recently, and are associated with contemporary pandemic outbreaks (W. Li et al. 2005; Vijaykrishna et al. 2007; Andersen et al. 2020; Boni et al. 2020; Banerjee et al. 2021).

Following its first detection in 2002 (Peiris et al. 2003), SARS-CoV-1 spread to more than 20 countries in six months, resulting in a short-lived but severe outbreak characterized by sustained human-to-human transmission (Cheng et al. 2007). Even though SARS-CoV-1 circulation was successfully halted, the virus underwent adaption to the human population, resulting in enhanced transmissibility (He et al. 2004). Since its emergence in 2019 (Zhou et al. 2020), SARS-CoV-2 has displayed highly efficient human-to-human transmission resulting in global spread, despite an apparently low rate of adaptive change during the early stages of the pandemic (van Dorp et al. 2020; MacLean et al. 2021). The constant circulation of OC43 and HKU1 in humans has been accompanied by continuous host-specific adaptation, a process now evident for SARS-CoV-2, represented by the ongoing emergence of virus lineages across time and space (with sub-lineages now reflecting regional ‘endemic’ patterns) (O’Toole et al. 2021). Thus, as SARS-CoV-2 becomes established in humans (Shaman and Galanti 2020), it will continue to adapt to overcome the selective pressures exerted by the collective immune response of the human host population (Kissler et al. 2020).

We hypothesize that adaptive convergence may occur across distantly related betacoronaviruses sharing the same ecological niche (in this case represented by the human host environment) (Gutierrez, Escalera-Zamudio, and Oliver G Pybus 2019). To test this hypothesis, we undertook a comparative analysis to search for evidence of shared mutational pathways between emerging sarbecoviruses that display sustained transmission in humans (particularly, SARS-CoV-2) and established human-endemic embecoviruses. Although MERS-CoV (*Merbecovirus* subgenus) can also infect humans, outbreaks have been the result of recurrent zoonotic events characterized by limited transmission chains, with no further evidence of adaptation (Fehr et al. 2017; WHO 2021). Thus, MERS-CoV and related viruses were excluded from our analysis.

Our approach involved the development of a methodological pipeline that allows the identification of amino acid changes putatively associated with adaptive convergence across multiple virus species. This is done by detecting non-synonymous mutations shared across virus taxa that display putative evidence of homoplasy (*i*.*e*., repeated mutations that do not share direct common ancestry) and/or stepwise evolution (*i*.*e*., sequential mutations leading towards a novel genotype) (for details see Supplementary Text 1 and Supplementary Figure 1). In parallel, we assess the impact of positive selection acting upon specific sites and/or branches (using dN/dS estimating methods), and determine the proximity of the mutations identified to viral protein surfaces with established function. Under these circumstances, structural contextualization of mutations may provide complementary information (to that derived from phylogenetic approaches), helping reveal the potential functional relevance for sites inferred as associated with adaptation across distinct virus species (Ellegren 2008; Hulswit et al. 2016; Avanzato et al. 2019; Escalera-Zamudio et al. 2020).

We identify 30 non-synonymous mutations shared across betacoronaviruses capable of sustained human-to-human transmission that display evolutionary patterns denoting putative homoplasy and/or stepwise evolution. The mutations identified were further ranked according to their selective relevance, and to their proximity to protein regions of known function. Under this approach, we subsequently identify four mutations (sites 18121 [nsp14/27], 21623 [spike/21], 21635 [spike/25] and 23948 [spike/796], in SARS-CoV-2 genome coordinates) near the functional surfaces of nsp14 (Ma et al. 2015), the spike protein S1 receptor-interacting subunit, and the S2 fusion machinery subunit. Our results provide a molecular-level context for common evolutionary trajectories that betacoronaviruses may undergo during their adaptation to the human host.

## RESULTS

### Patterns of genetic variability observed in human-infecting betacoronaviruses

Based on an alignment of the Orf1ab and S viral genes comprising approximately 1400 sequences (Methods section 1), we reconstructed the genomic diversity of the four distantly related betacoronavirus species studied here (Methods section 2). The patterns we recover following phylogenetic inference (represented by the expanded tree shown in Figure 1) are consistent with previously published phylogenies of the genus (Woo et al. 2006; Woo et al. 2010; Oong et al. 2017; Zhu et al. 2018; Bedford 2021). In this context, virus sequences belonging to the *Embecovirus* (HKU1, OC43, and related viruses) and the *Sarbecovirus* (SARS-CoV-1, SARS-CoV-2, and related viruses) (ICTV et al. 2017) subgenera form four well-supported clades. We were further able to identify all previously described genotypes for HKU1 (A-C) and OC43 (A-H) (Woo et al. 2006; Oong et al. 2017) (Supplementary Data 1). In summary, derived from genomic diversity reconstructions, our observations validate the evolutionary relationship known for these four distantly related betacoronaviruses.

**Figure 1.**
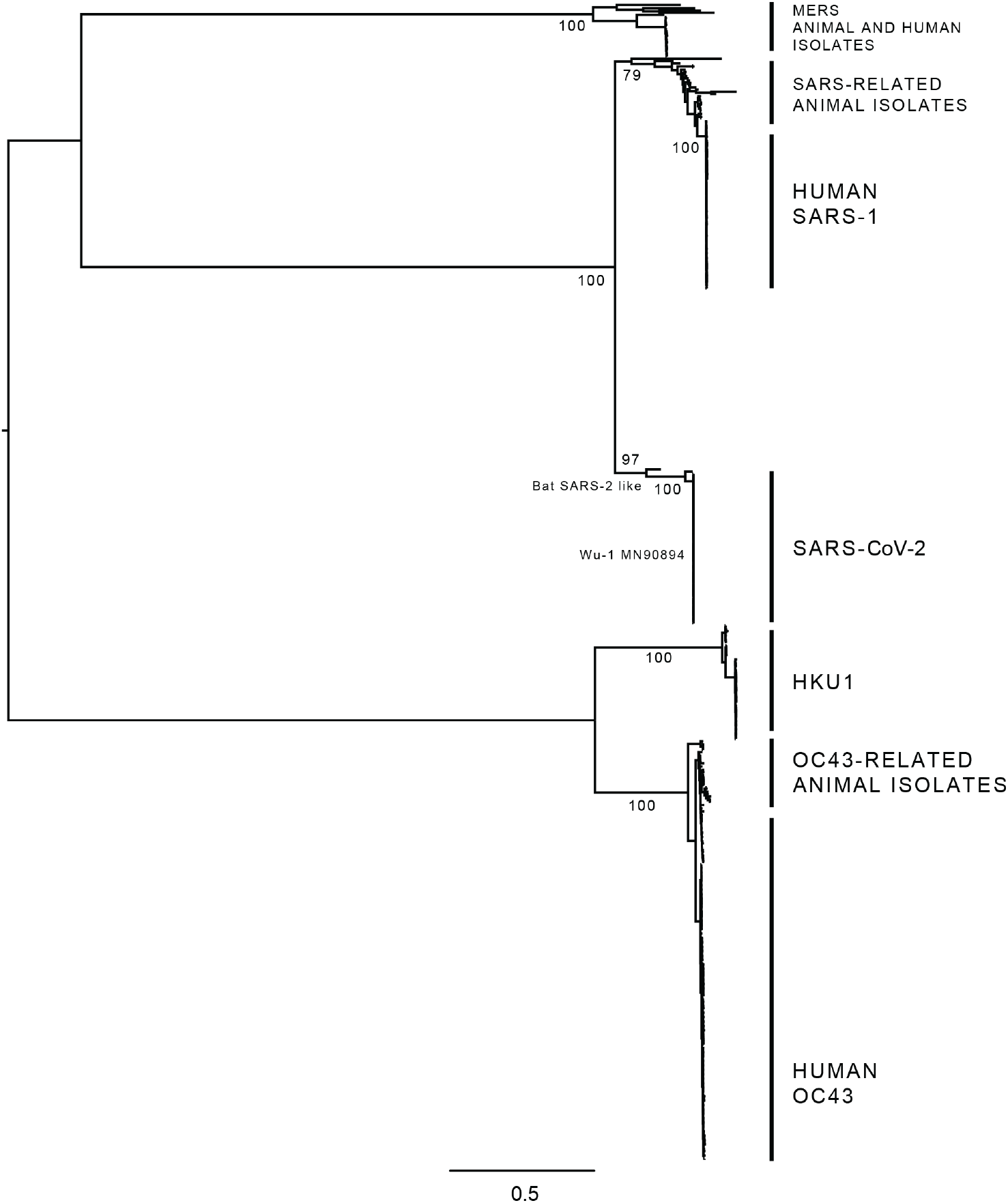
Phylogenetic tree of human-infecting betacoronaviruses. The expanded tree estimated from the Orf1ab+S alignment comprising 1455 sequences (see Methods section 6), summarizing the phylogenetic pattern observed for four distantly related human-infecting betacoronaviruses: HKU1, OC43, SARS-CoV-1 and SARS-CoV-2. MERS and related virus sequences were included in the tree for rooting purposes only. Both the *Embecovirus* subgenus (HKU1 and OC43 and related viruses) and the *Sarbecovirus* subgenus (SARS-CoV-1 and SARS-CoV-2 and related viruses) are indicated, showing the positioning of the most closely related virus genome sequences derived from animal isolates (when available). The different genotypes identified for the HKU1 (A, B and C) and for the OC43 (A–H) are shown in Supplementary Data 3.

Approximately 2% of the sites within the Orf1ab+S alignment (205/8962 codons) correspond to non-synonymous mutations shared between embeco- and sarbecovirus species (*i*.*e*., those present in any of the sarbecovirus clades, and also in HKU1 and/or OC43). For Orf1a, 2.7% of all sites (129/4774 codons) were identified as shared, whilst for Orf1b, only 0.9% of all sites (25/2623 codons) were shared. The highest proportion of shared mutations (3.2% of all sites, 48/1457 codons) was identified within Orf S. When analysing genetic variation within single virus species, a high degree of conservation was observed across Orf S (with >91% identity observed for all species). In general, most conserved sites we detect mapped to the membrane proximal S2 domain, whilst most variable sites mapped to the membrane distal S1 subunit (Figure 2). The predominance of variable sites within S1 compared to S2 was most evident for embecoviruses, and less so for sarbecoviruses; thus, we speculate that this may reflect a differential adaptation stage for the sarbecoviruses to the human host environment, evidenced by a lower degree of genetic divergence observed within Orf S.

**Figure 2.**
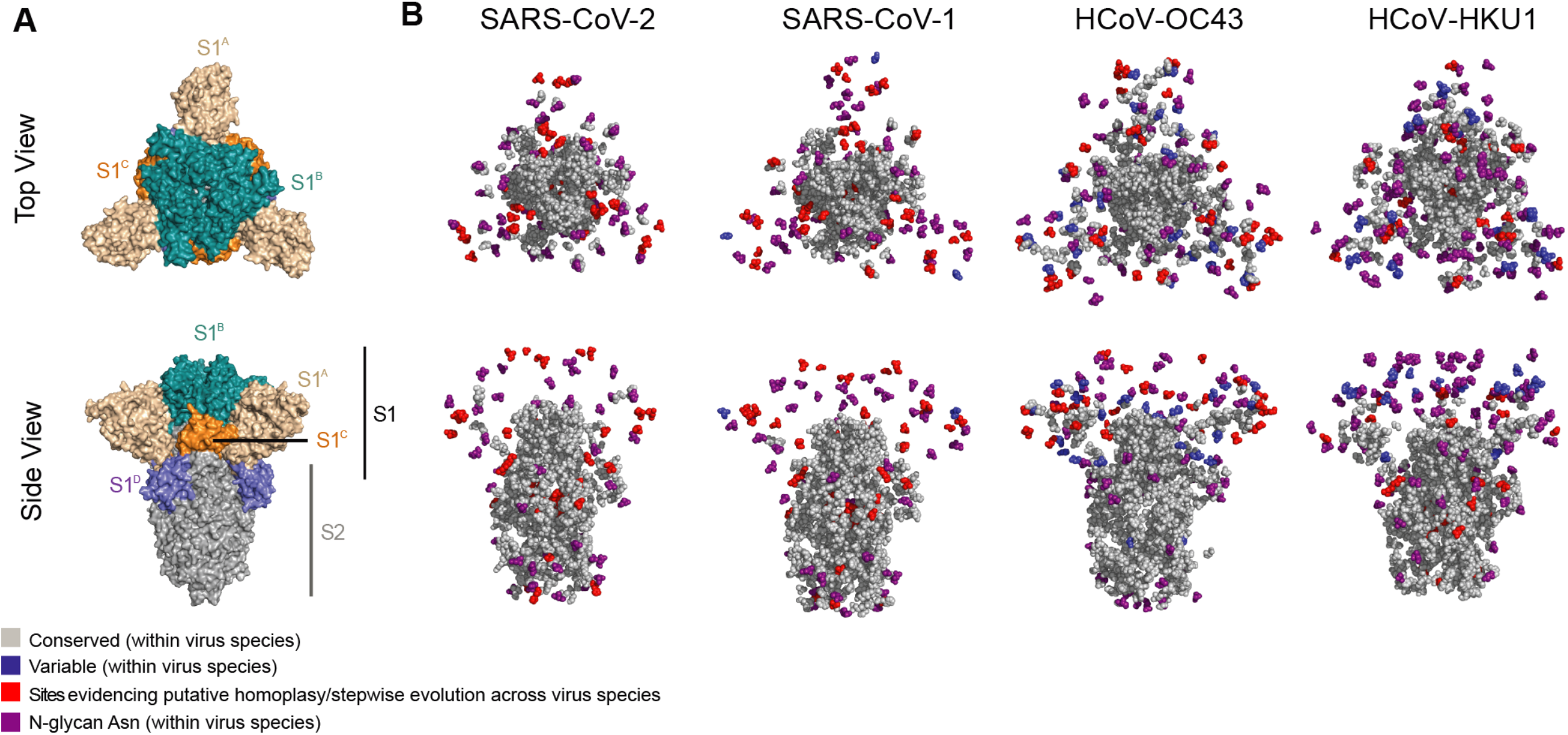
Distribution of conserved/variable sites with S across different virus species. (a) Top-down (upper panel) and side view (bottom panel) of a cartoon representation of the multidomain architecture of the trimeric SARS-CoV-2 S ectodomain (PDB: 6VXX). The S1 subunit is coloured according to the different protein domains: S1^A^ in cream, S1^B^ in teal, S1^C^ in orange, and S1^D^ in blue, whilst the S2 subunit is shown in grey. (b) Top-down and side views of sphere-based representations of trimeric S protein ectodomains for the viruses studied here: SARS-CoV-2 (PDB: 6VXX), SARS-CoV-1 (PDB: 6ACC), OC43 (PDB: 6OHW) and HKU1 (PDB: 5I08). The sphere-based representation shows conserved (shown in grey; residues present ≥99% of all sequences) and variable sites (blue; residues present in ≥1% of all sequences) across virus species. Variable sites identified as denoting homoplasy or stepwise evolutionary patterns are shown in red (see Methods section 3). The asparagine residues of N-linked glycosylation sequons are indicated in purple.

When analysing genetic variation across virus species, the Orf S genomic region displayed a greater proportion of variable sites relative to the number conserved sites (for the definitions used for ‘conserved’ and ‘variable’ sites, see Methods section 3), as has been before noted for other coronaviruses (Hulswit et al. 2016). In this context, only 16% of the homologous sites within the Orf S alignment were found to be conserved, whilst the remaining 84% were variable (Supplementary Data 2). The highest proportion of conserved sites locate to the S2 subunit, presumably reflecting functional constraints of the viral membrane fusion machinery shared across coronavirus species (Li 2016). Within the S1 subunit, a greater number of variable sites were observed, particularly within the S1^A^ domain (also known as the N-terminal domain, or NTD). In contrast, the S1^B^ domain did not display any conserved sites across virus species, likely associated with differences in receptor engagement process between embeco- and sarbecoviruses (mediated by the S1 subunit), as embecoviruses use the S1^A^ domain to interact with sialoglycan-based receptors, whilst sarbecoviruses use their S1^B^ domain to bind to angiotensin-converting enzyme 2 (ACE2) (Hulswit et al. 2019; Lan et al. 2020). Additionally, a conserved R residue at site 685 within the S1/S2 cleavage site (numbering according to the SARS-CoV-2 protein, codon sites 23615-23617) was found to be shared within and across all virus species (Supplementary Data 2), reflecting the conserved proteolytic maturation mechanism of the spike protein across virus taxa belonging to the *Orthocoronavirinae* family (Millet and Whittaker 2015).

### Sites displaying evidence of homoplasy and/or stepwise evolution

Although not all non-synonymous mutations putatively displaying homoplasy and/or stepwise evolution may derive from positive section, such mutational patterns are most likely to result from adaption (Stern et al. 2017; Gutierrez, Escalera-Zamudio, and Oliver G. Pybus 2019; Escalera-Zamudio et al. 2020). Thus, amongst the non-synonymous mutations identified as shared across virus species, we further searched for those displaying putative evidence for homoplasy and/or stepwise evolution (Supplementary Text 1) under our pipeline (Methods section 3). After visual validation, we confirmed that 30 sites (representing 0.3% within the Orf1ab+S alignment) display evolutionary patterns indicative of homoplasy and/or stepwise evolution (see Supplementary Text 2, Supplementary Figure 2 and 3). Two of these were found within Orf1a, nine within Orf1b, and 19 within Orf S (Table 1). The evolutionary trajectories of different amino acid states observed for three illustrative sites (18121, 21623 and 23948) are highlighted below and in Figure 3.

**Table 1.**
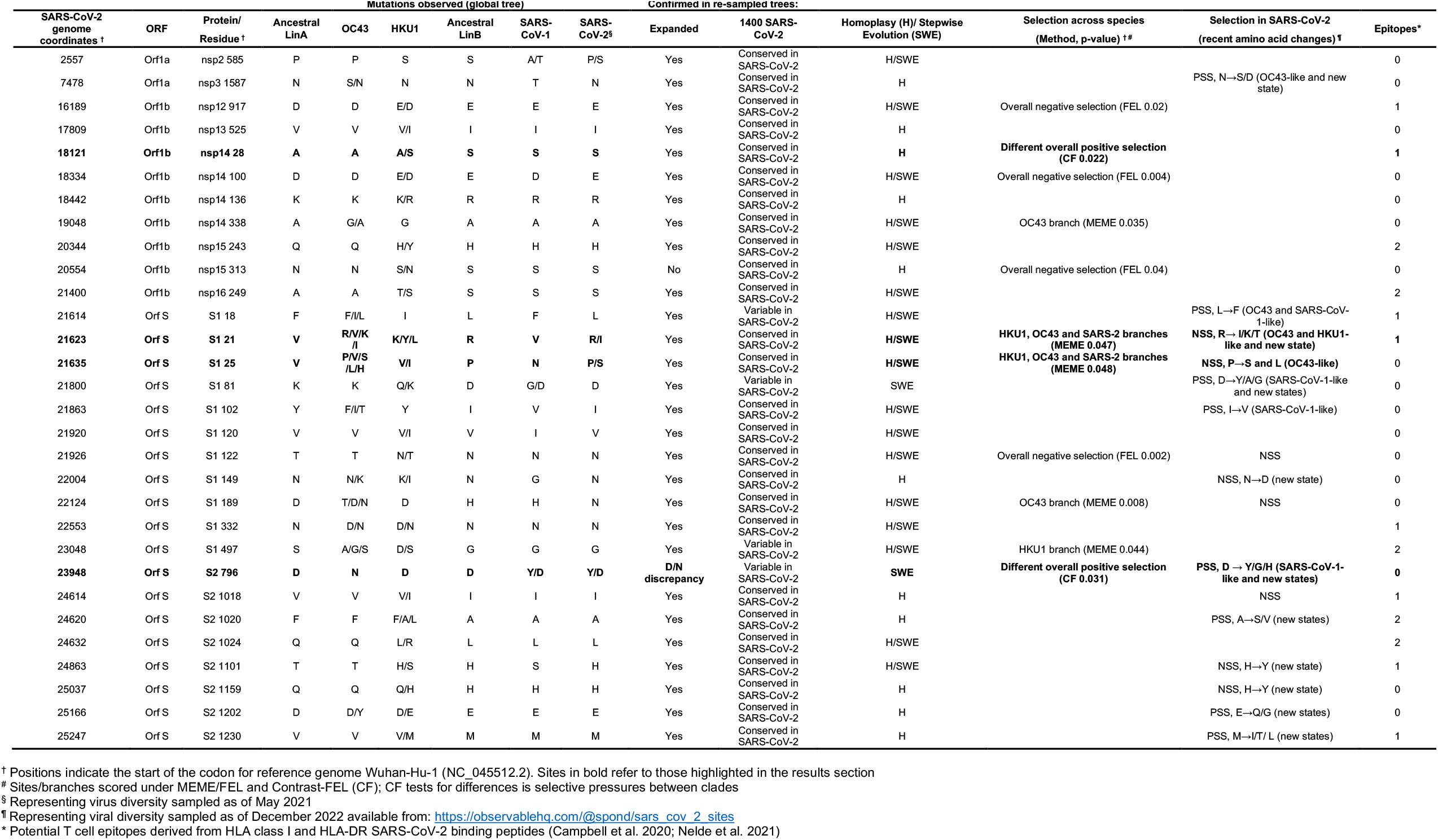
Potentially relevant sites across human-infecting betacoronaviruses.

**Figure 3.**
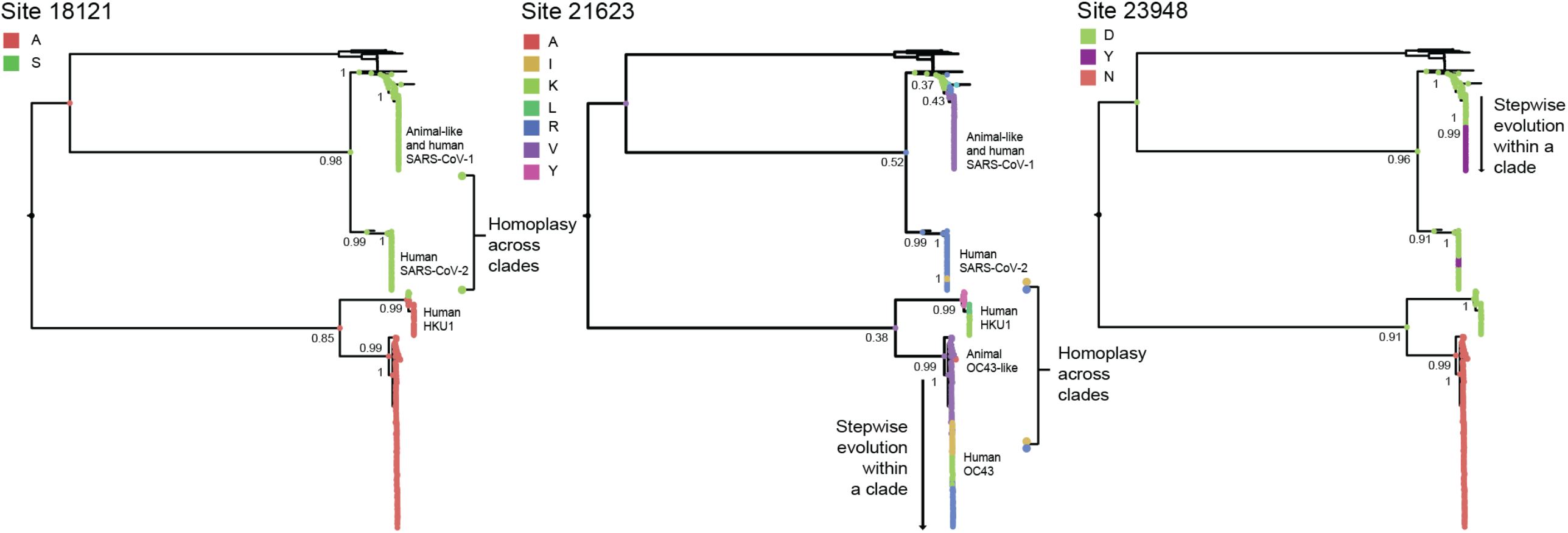
Reconstruction of amino acid evolution at selected sites. Maximum clade credibility (MCC) trees showing the evolutionary trajectories for different amino acid states observed for three illustrative sites (18121, 21623 and 23948) that (i) display evidence of homoplasy and/or stepwise evolution, (ii) show evidence of positive selection, and (iii) are proximal to regions of established protein function. The reconstructions of ancestral states for these sites show different amino acid states at nodes (represented by circles in different colours). The posterior probability for a given amino acid state occurring at a given node of interest is indicated. Sites 18121 display evidence of homoplasy across virus lineages, site 21623 shows evidence of both homoplasy across species and stepwise evolution within single virus species (*i*.*e*. OC43), and site 23948 shows evidence of stepwise evolution within single virus species (*i*.*e*. SARS-CoV-1), and also of homoplasy across virus species (*i*.*e*. SARS-1 and SARS-CoV-2).

Derived from the global, expanded and the re-sampled SARS-CoV-2 trees (Methods section 1, 3 and 6, Supplementary Text 3), our results show that site 18121 (codon 18121-18123 in Orf1b/nsp14, corresponding to amino acid state ‘S’) is homoplasic between HKU1 genotype B and the sarbecoviruses (Table 1, Figure 3, Supplementary Data 3). Comparably, site 21623 (codon 21623-21626 in Orf S, corresponding to amino acid state ‘R’) was identified as homoplasic between SARS-CoV-2 and OC43 genotypes D, F, G and H. This site also displayed evidence for stepwise evolution within a single virus clade (OC43), exemplified by the sequential amino acid replacement pattern of V⟶ I⟶ K⟶ R (Figure 3).

For site 23948 (codon 23948-23950 in Orf S), initial observations based on the global tree revealed that amino acid state ‘D’ was present in all virus species, except for OC43 (displaying amino acid state ‘N’). However, when replicating our analyses (expanded tree), the distribution of amino acid state ‘D’ was now present in some embecoviruses (including OC43 but excluding HKU1) and most sarbecoviruses. These discrepancies are likely due to alignment uncertainty across genome regions of highly divergent virus taxa. Nonetheless, based on consensus protein sequences and structural comparison, the structural contextualization of 23948 and adjacent sites confirmed the presence of ‘D’ in SARS-CoV-1, SARS-CoV-2 and HKU1, and ‘N’ in OC43, (Supplementary Figure 6). Thus, amino acid state ‘D’ at site 23948 shows evidence of homoplasy between the SARS-CoV-1, SARS-CoV-2 and HKU1.

For this same site (23948), an additional amino acid change from ‘D’ to ‘Y’ was identified as homoplasic between some SARS-CoV-1 and SARS-CoV-2 sequences (data derived from the global, expanded and the re-sampled SARS-CoV-2 trees) (Table 1). For SARS-CoV-2, amino acid state ‘Y’ emerged and was lost repeatedly during the early stage of the pandemic (represented by independent minor clusters that became extinct). However, following emergence and global spread of the B.1.1.529 virus lineage (Omicron variant of concern [VOC], and descending sub-lineages), amino acid state ‘Y’ replaced amino acid state ‘D’, displaying a prevailing trend associated with the dominance of the B.1.617.2 lineage (Delta VOC, and descending sub-lineages) (Table 1, Figure 3, Supplementary Data 3) (also confirmed by independently sampled SARS-CoV-2 data available up to December 2022: https://nextstrain.org/groups/neherlab/ncov/global?c=gt-S_796).

### Quantifying the effects of positive selection

The dN/dS estimates we obtained across complete virus genomes and upon specific coding regions (see Methods section 4) indicates that positive selection is acting upon the Orf1ab and Orf S of SARS-CoV-2, compared to other viruses studied here. Specifically, the effect of episodic diversifying selection was detected upon 5/14 non-recombinant fragments (three in Orf1b and two in Orf S, for details see https://observablehq.com/@spond/beta-cov-analysis). Using the Contrast-FEL method to detect the effect of a differential selection across branches separating lineages (see Methods section 4), we found 36 sites (0.4%) evolving under differential selective pressure across distinct virus clades. Furthermore, we found 0.7% of all sites (67 codons within the Orf1ab+S alignment) to be evolving under episodic diversifying positive selection (scored under MEME with a p≤0.05 as *positively-selected sites*, or PSS) (Supplementary Table 2). In contrast, we found 5% of all sites (461 codons within the Orf1ab+S alignment) to be evolving under pervasive negative selection (scored under FEL with a p≤0.05 as *negatively-selected sites*, or NSS). We subsequently mapped the identified PSS and NSSs onto the SARS-CoV-2 S protein structure (Methods section 5). We observe that out of a total of 22 PSSs, 18 locate within the S1 subunit (11 in S1^A^, 5 in S1^B^, 1 in S1^C^ and 1 in S1^D^ domains), whilst the remaining four mapped onto the S2 subunit. Conversely, out of a total of 82 of NSSs, 46 locate within S1 (18 in S1^A^, 21 in S1^B^, 3 in S1^C^ and 4 in S1^D^), whilst the remaining 36 mapped onto S2 (Supplementary Figure 7).

From the 30 non-synonymous mutations we identify as displaying evolutionary patterns putatively denoting homoplasy and/or stepwise evolution (Table 1), sites 19048, 21623, 21635, 22124 and 23048 were further scored as PSS (under different methods). Sites 21623 and 21635 were inferred as PSSs along ancestral branches leading to the HKU1, OC43 and SARS-CoV-2 clades. Sites 19048 and 22124 were inferred as PSSs along the OC43 ancestral branch, whilst 23048 was inferred as a PSS along the HKU1 ancestral branch (Table 1, Supplementary Table 2). Further analysis under the branch and site model in the MEME method (Methods section 4) revealed site 18121 to be evolving under positive selection for the HKU1 clade/branch (relative to the sarbecoviruses), in agreement with our observations made on putative homoplasy detected for this site between HKU1 genotype B, SARS-CoV-1 and SARS-CoV-2 (Table 1, Figure 3). Similarly, site 23948 was also inferred to be evolving under positive selective for the SARS-CoV-1 branch, relative to other virus clades (Supplementary Table 2).

For validation purposes, we compared our results with selection analysis available for independently sampled SARS-CoV-2 genome data available as of December 2022 (https://observablehq.com/@spond/evolutionary-annotation-of-sars-cov-2-covid-19-genomes-enab) (Kosakovsky Pond). Of the 30 mutations we identify, 16 of these are currently scored as PSS or NSSs, with 13 of these mapping directly onto potential T cell epitopes derived from HLA class I and HLA-DR binding peptides in SARS-CoV-2 (Campbell et al. 2020; Nelde et al. 2021) (Table 1). Additionality, up to December 2022, sites 7478, 21614, 23948, 24620 and 25166 were detected as evolving under positive selection, whilst sites 21635, 24863, and 25037 were detected as evolving under negative selection.

### Contextualization of mutations using protein structural and functional information

For this purpose, we mapped the 30 mutations identified (Table 1) onto corresponding protein structures (Methods section 5). Below, we focus on the potential functional impact identified for four sites (18121, 21623, 21635 and 23948) that meet three criteria of: displaying evidence of homoplasy and/or stepwise evolution, showing evidence of evolution under positive selection, and being proximal to regions of established protein function. A description for all other identified mutations is available in the Supplementary Text 4 and Supplementary Table 3.

#### Site 18121 in Orf1ab

The Orf1ab gene encodes 16 non-structural proteins (nsp1-16), primarily involved in viral RNA synthesis and processing (V’kovski et al. 2020). Site 18121 is located within Orf1ab and corresponds to an ‘S’ to ‘A’ mutation at residue 28 within the exonuclease domain of the nsp14 protein (numbering according to the SARS-CoV-1 protein) (Supplementary Table 3, Figure 4). Nsp14 functions as a methyltransferase, and is involved in 5′-capping of the viral mRNA (Ma et al. 2015). The cap core structure is essential for viral mRNA transcription, but is also implicated in protecting the 5′-triphosphate from activating the host innate immune response (Wang et al. 2015). An ‘S’ to ‘A’ mutation within this region is expected to result in the loss of an intra-protein hydrogen-bond (formed with the main chain of residue T25 of nsp14, and between nsp14 and its activator nsp10) (Ma et al. 2015), potentially modulating protein-protein interaction (Figure 4) (assessed under PISAebi; http://www.ebi.ac.uk/pdbe/prot_int/pistart.html) (Krissinel and Henrick 2007).

**Figure 4.**
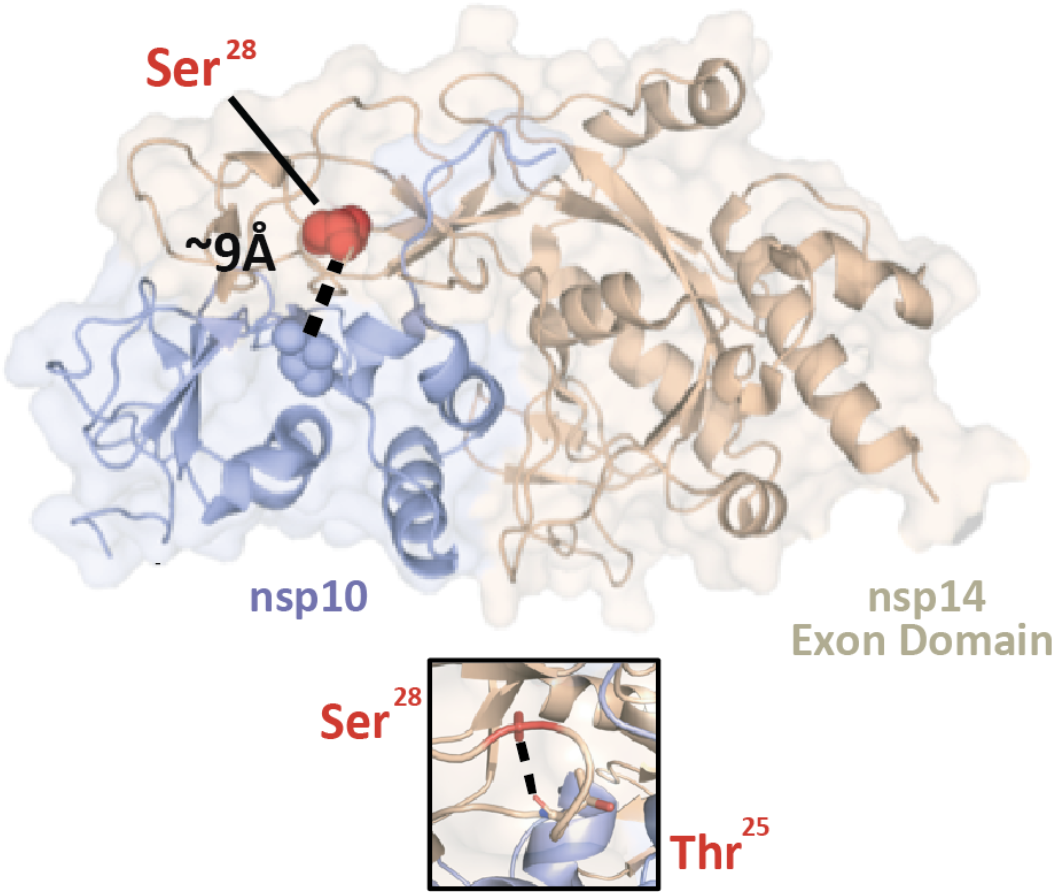
Residue Ser^28^ of nsp14 is situated near the nsp14-nsp10 interface. Cartoon representation of the SARS-CoV-1 nsp14-nsp10 protein complex (PDB: 5C8S) with Ser^28^ (corresponding to site 18121 in SARS-CoV-2 genome coordinates) shown as a red sphere. This residue is located within the nsp14 ExoN domain (cream) and is approximately 9 Å from the interface with nsp10 (light blue, the proximal nsp10 residue Cys^41^ was used to calculate the distance and is indicated as a sphere). The distance between nsp14’s Ser^28^ and nsp10’s Cys^41^ is annotated and indicated by a dashed black line. Zoomed-in panel: detailed representation of the intra-nsp14 hydrogen-bond between the side chain of Ser^28^ and the main chain of Thr^25^ (identified with the PISAebi server). The side chain of Ser^25^ is indicated as a red stick and Thr^25^ is indicated in sticks and coloured according to atom (C, cream; O, red; N, blue). The hydrogen-bond is indicated as a dashed black line.

#### Sites 21623 and 21635 in S1

The S1 subunit of S (spike) protein mediates attachment of the virus to the host cell (Li 2016). Humans-infecting embecoviruses bind to glycan-based cell receptors terminating in 9-*O*-acetylated sialic acids, and receptor recognition is accomplished via two hydrophobic pockets separated by a conserved ‘W’ residue within the S1^A^ region of the protein (Hulswit et al. 2019; Tortorici et al. 2019). In contrast, the receptor-binding site for human-infecting sarbecoviruses consists of an extended loop located within the S1^B^ domain of the protein (Li et al. 2005; Lan et al. 2020; Shang et al. 2020). Despite a limited degree of conservation amongst contact residues within the RBD of SARS-CoV-1 and SARS-CoV-2, both virus species recognize the ACE2 molecule to enter the host cell (Li et al. 2005; Lan et al. 2020).

Site 21623 displays several non-synonymous mutations (‘R’, ‘V’, ‘K’ and ‘I’) mapping to residue 29 of the OC43 S protein, located within the S1^A^ domain of the S1 subunit (Supplementary Table 3, Figure 5). Site 21635 again shows multiple non-synonymous mutations ‘P’, ‘V’, ‘S’, ‘L’ and ‘H’ mapping to residue 33 of the OC43 S protein. Residue 29 (site 21635) is just four positions downstream of residue 33 (site 21623) (Figure 5). Within the OC43 S protein, both residues 29 and 33 are located within a loop neighbouring the hydrophobic pockets in S1^A^. This area is instrumental for receptor recognition, and changes within the region can modulate receptor affinity (Hulswit et al. 2019). Our observations on both these sites displaying mutational patters putatively denoting homoplasy/stepwise evolution and evidencing positive selection (Table 1), are congruent with antigenic drift shaping the evolution of human-endemic coronaviruses (Kistler and Bedford 2021).

**Figure 5.**
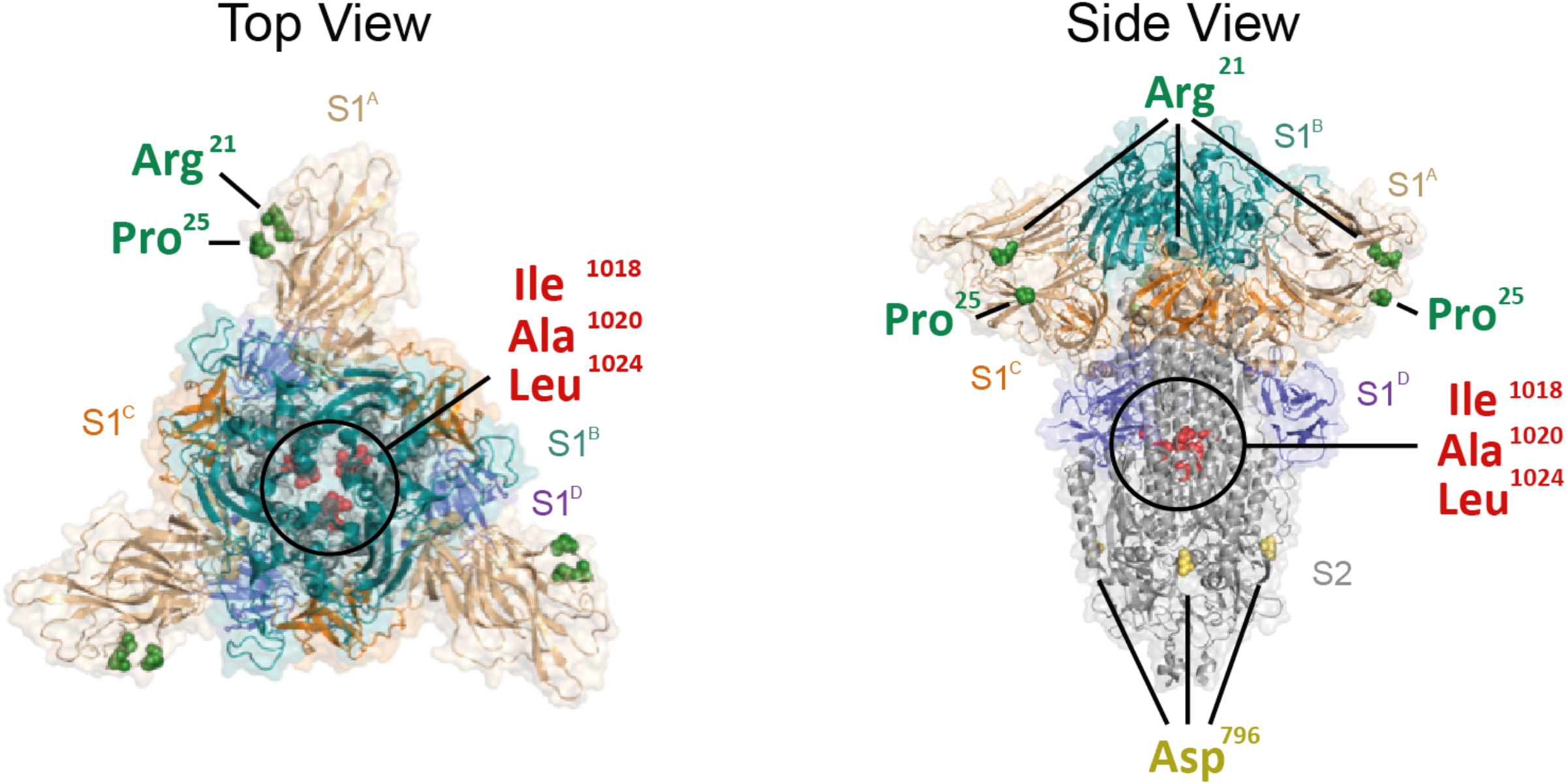
Mapping of mutations exhibiting homoplasy onto the S protein structure of SARS-CoV-2. Top-down (left) and side view (right) of a cartoon representation of the multidomain architecture of the trimeric SARS-CoV-2 S ectodomain (PDB: 6VXX). The S2 subunit is highlighted in grey and the S1 ectodomain is divided into S1^A^ (highlighted in cream), S1^B^ (teal), S1^C^ (orange), and S1^D^ (blue) domains, following the colour scheme in Figure 3. Homoplasic mutations that co-localize to known functional surfaces (see Supplementary Table 4) are indicated in the structure and coloured in groups: Arg^21^ (corresponding to site 21623 in SARS-CoV-2 genome coordinates, in green), Pro^25^ (site 21635, in green), Asp^796^ (site 23948, in yellow), Ile^1018^ (site 24614, in red), Ala^1020^ (site 24620, in red) and Leu^1024^ (site 24632, in red). All representations are shown with a transparent protein surface for clarity.

As a note, in the case of SARS-CoV-2, mutations in both these sites have been observed for two VOCs (B.1.351 and P.1, ‘Beta’ and ‘Gamma’) (Faria et al. 2021; Tegally et al. 2021). Particularly, site 21623 corresponds to variable non-synonymous mutations (‘R’, ‘I’, ‘T, ‘K and ‘G’) mapping to residue 21 in the S protein (as confirmed by independently sampled SARS-CoV-2 genome data available up to December 2022: https://nextstrain.org/groups/neherlab/ncov/global?c=gt-S_21). However, given that sarbecoviruses engage with the ACE2 receptor via domain S1^B^, mutations observed within domain S1^A^ may reflect relaxed constraints in the local protein surface, unlikely to be related to the receptor’s functionality.

#### Site 23948 in S2

Virus binding to the cell surface is followed by fusion of the viral envelope with the host cell membrane, thereby delivering the viral genome into the target cell. The S2 subunit harbours the viral fusion machinery and displays characteristic structural features of class I fusion proteins. This includes a fusion peptide that anchors the virus to the host membrane, initiating fusogenic conformational changes that result in merging of the two membranes (Li 2016). However, for this process to occur, the S protein needs to be primed through cleavage initiated by host cell proteases (Millet and Whittaker 2015). In coronaviruses, cleavage occurs at a conserved site referred to as the S1-S2 junction (consensus RRAR|S in SARS-CoV-2), and is further mediated by an additional R|S site located immediately upstream of the fusion peptide in the S2 subunit (referred to as the S2’ cleavage site) (Millet and Whittaker 2015).

For site 23948, the observed mutation ‘D’ to ‘Y’ (identified as homoplasic between some SARS-CoV-1 and SARS-CoV-2 sequences) corresponds to a non-conservative amino acid replacement at residue 796 of the S2 subunit (SARS-CoV-2 numbering) (Table 1, Supplementary Table 3). This residue is located immediately upstream of the S2’ cleavage site, within the region that crucially regulates the release of the fusion peptide. Residues located between position 796 and the fusion peptide form a loop, displaying variability across betacoronavirus species (Supplementary Figure 6), suggesting that the apparent relaxed local constraints may serve to secure loop accessibility for cleavage activation. In agreement with this hypothesis, the corresponding protein region within the HKU1 structure remains unresolved (Kirchdoerfer et al. 2016).

## DISCUSSION

In this study, we searched for signatures of adaptive convergence across distantly related human-infecting betacoronaviruses, represented by shared non-synonymous mutations that putatively denote homoplasy and/or stepwise evolution. The mutations we identified were further ranked according to their selective relevance, and to their proximity to protein regions of known function. The majority of the mutations we observe locate to the receptor binding region of the S protein (*i*.*e*., S1 subunit), whilst a smaller proportion of these were found within non-structural proteins encoded by Orf1ab (site 18121 in the exonuclease domain of nsp14, and site 20344 in the endonuclease domain of nsp15). Further *in-silico* analyses revealed four genomic sites (18121, 21623, 21635 and 23948) that display cumulative evidence of: *i*) a mutational pattern putatively denoting homoplasy and/or stepwise evolution, *ii*) evolution under positive selection, and *iii*) being structurally proximal to regions of known protein function. Below, we discuss our findings in light of three key evolutionary processes: antigenic drift, epistasis and adaptive convergence.

The host humoral immune response is an evolutionary force driving antigenic drift in viruses. In the case of betacoronaviruses, this is reflected by cumulative mutations in the S protein (particularly within the S1 subunit) that may allow frequent reinfections of the host population (Forni et al. 2021; Kistler and Bedford 2021; Yewdell 2021). In agreement with this observation, the emergence of some SARS-CoV-2 lineages (particularly VOC) has been associated with high levels of infection in pre-exposed human populations across different geographic regions (as an example on P.1, see Faria et al. 2021). Our results evidence antigenic drift upon the S1 subunit of distinct betacoronaviruses as a major component of the adaptation process to the human host environment, further evidenced by Orf S also being the least conserved genome region across distinct virus species (Li 2016). On the other hand, mutations found within Orf1ab could have a potential impact on viral fitness related to an enhanced replication efficacy in the human host (Menachery et al. 2017). As the evolution of Orf1ab is also driven by immune responses such as cytokine signalling cascades and antigen presentation (Wang et al. 2015; Hackbart et al. 2020; Taefehshokr et al. 2020; Yuen et al. 2020), these mutations may also be the result of concerted selective pressure(s), following the principle that single mutational changes can have pleiotropic effects on distinct viral phenotypes and fitness components (de Wilde et al. 2018).

Identifying adaptive convergence raises the possibility of predicting mutational pathways in viruses relevant to global health (Gutierrez, Escalera-Zamudio, and Pybus 2019). When applied to SARS-CoV-2, our results reveal that some of the mutations we had initially identified as potentially relevant back in May 2021 (see (Escalera-Zamudio et al. 2021) had already been observed in other betacoronaviruses that circulate endemically in humans (Table 1), and some now display dominant trends in SARS-CoV-2 (as analysed up to December 2022). For example, amino acid state ‘R’ at residue 21 of the S protein (sites 21623) (https://nextstrain.org/groups/neherlab/ncov/global?c=gt-S_21) and ‘P’ at residue 25 (site 21635) (https://nextstrain.org/groups/neherlab/ncov/global?c=gt-S_25) have dominated across time. Moreover, mutation ‘D’ to ‘Y’ observed at residue 796 of the S protein (site 23948) has proven to be a successful mutational pathway, evidenced the replacement of state ‘D’ (previously observed for the B.1.617.2 lineage, Delta VOC and descending sub-lineages) by state ‘Y’ (now observed for the B.1.1.529 lineage, Omicron VOC and descending sub-lineages) (https://nextstrain.org/groups/neherlab/ncov/global?c=gt-S_796). Of interest, mutations at residue 796 of the S protein have been linked to the emergence viral variants that display reduced susceptibility to neutralizing antibodies (Kemp et al. 2021).

Epistasis is thought to have played a central role in the emergence of human-infecting betacoronaviruses (Holmes and Rambaut 2004). However, inferring epistasis across diverging viruses is difficult given the functional differences between homologous genes and proteins. Through our methodological approach we cannot measure epistasis *per se*, but we can aim to identify adaptive convergence and subsequently discuss its possible effects in the light of epistasis. Thus, our results indirectly provide support for epistasis, in the sense that if the same amino acid changes are observed in different virus species, then associated epistatic interactions are expected to be shared. This is of particular importance when considering the potential role of epistasis in antigenic drift, where the combined effect of independent mutations could contribute to antigenic escape (Rochman et al 2022). In the context of our findings, sites 21623 and 21635 are presumed to be involved in the antigenic drift of embecoviruses. As these residues are in close proximity to each other (displaying a linked evolution), these could thus reflect epistatic interactions. Nevertheless, within the SARS-CoV-2 S1^B^-ACE2 interface, epistasis seems to play a limited role, as the effect of multiple mutations seems to be additive rather than epistatic (Zahradník et al. 2021; Rochman et al. 2022; Starr et al. 2022).

The mutational spectrum of SARS-CoV-2 is known to be impacted by the human host apolipoprotein B mRNA-editing enzyme (APOBEC) family (Di Giorgio et al. 2020). The activity of APOBEC induces C ⟶ U/T mutations in the viral genome through a cytidine deaminase activity, likely resulting in a high degree of apparent homoplasy, reflected in emerging mutations across distinct virus sub-populations (De Maio, et al. 2020; Worobey, et al. 2020; Wang, et al. 2021). Relative to more commonly used strategies for identifying homoplasy within single virus species, our methodology poses an alternative approach that aims to identify homoplasy *across and within* virus taxa, as represented by shared mutations fixed under an evolutionary scenario mostly driven by selection (see Supplementary Text 5). Given that candidate mutations are observed over longer evolutionary times, this approach represents a useful tool that decreases the likelihood of erroneously scoring mutations as homoplasic (such as those resulting from mutational biases inherent to the SARS-CoV-2 genome).

However, identifying adaptive convergence faces several important limitations. First, the methodology we use is conservative, as it is based on strict homology. In this context, we only consider sites robustly identifiable as homologous that can be traced back to ancestral nodes with confidence (consequently excluding highly divergent genes). Therefore, our approach may result in an underestimation of sites that may putatively denote adaptive convergence across divergent virus species. Moreover, a limited virus genome sampling across time and space (in particular for HKU1 and SARS-CoV-1), coupled with a relatively low genetic diversity observed for SARS-CoV-2 (Rausch et al. 2020), further restricts the potential to identify shared mutations across virus species (van Dorp et al. 2020). In addition, there is some uncertainty associated with mutations that may derive from processes such as founder effects, mutational hitchhiking, and toggling at hypervariable sites (Delport et al. 2008; Pond et al. 2012; Simmonds 2020; Wang et al. 2020; De Maio et al. 2021). Finally, whilst our analysis provides insights into coronavirus evolution in humans, our approach renders us unable to identify mutations that may result from host switching events. This is due to analyses on nodes representing ancient host switching events (Corman et al. 2018) being constrained by long divergence times, differences in mutation rates across virus taxa in different hosts, mutational saturation, and by a considerable under-sampling of betacoronaviruses circulating in non-human hosts (Holmes and Rambaut 2004; De Maio et al. 2021).

In this sense, additional/future experimental data could help reveal the impact of mutations on viral fitness. However, performing such studies may be difficult, as these concern potential gain-of-function experiments. Alternatively, enhanced genomic surveillance of betacoronaviruses infecting the human population and of those ciruclating in other animal host may confirm whether the mutational pathways we identify here represent evolutionary trajectories on which betacoronaviruses converge in their adaptation process to the human host.

## MATERIAL AND METHODS

### 1. Initial data collation

When this manuscript was first deposited as a preprint (May 2021) (Escalera-Zamudio et al. 2021), complete genomes for all HKU1, OC43 and SARS-CoV-1 viruses sampled across different geographical regions and time were downloaded from the Virus Pathogen Resource (ViPR-NCBI 2021) (Supplementary Data 4). Sequences were removed if meeting any of the following criteria: (i) being >1000nt shorter than full genome length, (ii) being identical to any other sequence, or (iii) if showing >10% of site were ambiguities (including N or X). A total of 53 HKU1, 136 OC43 and 40 SARS-CoV-1 sequences were initially retained for analyses. For SARS-CoV-2, to better reflect an early zoonotic process into the human population (MacLean et al. 2021), we originally aimed to limit the genetic diversity of the sampled virus population to the first wave of infection recorded during the pandemic. For this, ∼23000 full genomes sampled worldwide before May 2021 available in the GISAID platform (GISAID 2021) were downloaded and aligned as part of an initial public dataset provided by the COG-UK consortium (COG-UK 2021) (Supplementary Data 4). To make local analyses computationally feasible, the original SARS-CoV-2 dataset was randomly subsampled to ∼5% of its original size, keeping the earliest genomes, and further reducing the dataset under the quality criteria stated above. In total, 1120 SARS-CoV-2 sequences were retained. For all virus species considered in this study, we focused only on genomes derived from human cases, in order to reflect host-specific adaptation processes.

### 2. Phylogenetic analysis

Only the main viral ORFs (Orf1ab and S) were used for further phylogenetic analysis, as these are homologous amongst the four viral species studied, and also encode proteins that are essential for viral function (*i*.*e*., the genome replication machinery, receptor binding, and the virus-host membrane fusion apparatus) (Yoshimoto 2020). For each virus species, individual ORFs were extracted and aligned using MAFFT v7.471 (Katoh and Standley 2013), and were concatenated to generate Orf1ab+S alignments (Supplementary Data 1). The concatenated alignments were then combined to generate a global alignment dataset that was then re-aligned at amino acid level using a profile-to-profile approach following taxonomic relatedness (Wang and Dunbrack 2004). This global alignment comprised in total 1314 sequences and 26883 sites.

Maximum Likelihood phylogenies were estimated for the individual and global alignments using RAxML v8 (Stamatakis 2015), under a general time reversible nucleotide substitution model and a gamma-distributed among-site rate variation (GTR+G). Branch support was assessed using 100 bootstrap replicates. All trees were midpoint-rooted, whilst general phylogenetic patterns observed amongst these distantly related virus species were validated by comparing to previously published phylogenies (Woo et al. 2006; Woo et al. 2010; Lau et al. 2011; Oong et al. 2017; Zhu et al. 2018; Bedford 2021). Recombination is known to be common amongst betacoronaviruses (Woo et al. 2006; Su et al. 2016; Oong et al. 2017), including SARS-CoV-2 (Gutierrez et al. 2022; Turakhia et al. 2022). However, recombinant sequences were not removed at this step, as it was important to detect potentially recombinant isolates that could display relevant mutations. Putative recombinant sequences were eventually removed for subsequent analyses (when identified, see Methods sections 6 and 7).

### 3. Identifying homoplasy and/or stepwise evolution

Following the pipeline described by Escalera & Golden (Escalera-Zamudio et al. 2020), variable sites across different virus taxa were identified within the global alignment as those displaying non-synonymous mutations (rendering amino acid changes) occurring in at least ≥1% of the sampled sequences. Variable sites were extracted by masking columns across the alignment showing identical sites and at least 50% gaps, followed by the ‘Find Variations/SNPs’ function used to compare each site to a consensus generated under a 95% threshold with Geneious Prime v2020.0.4 (Kearse et al. 2012). A total of 6681 variable sites were identified and used to infer ancestral amino acid state reconstructions onto the nodes/internal branches of the global tree (see Methods section 2 above). This was done using TreeTime (Sagulenko et al. 2018) under a ML approach (RAS-ML) under a time-reversible model (GTR) for state transitions. The genetic variability observed within leaves/tips of the tree was deliberately excluded, in order to only analyse changes occurring within nodes or internal branches of the tree. In parallel, conserved sites were identified as those present in ≥ 99% of the sampled virus sequences. Conserved sites were extracted by reversing the ‘variable site masking’, to obtain only identical sites identified across the global alignment (Supplementary Data 2).

The resulting 6681 ‘Ancestral Reconstruction Trees’ (named here ARTs) were then classified under a computational algorithm developed to sort mutational patterns based on whether or not they support homoplasy and/or stepwise evolution. Briefly, homoplasy can occur within nodes of single clade or across clades, in which the same amino acid change must be present in at least one internal node of any given clade, and in another internal node of the same/another clade. Clades with the same amino acid states must not share direct common ancestry. Conversely, stepwise evolution is represented as sequential mutations occurring at the same sites within a single clade. Any given site scored under putative ‘stepwise evolution’ must display changes between at least two different states (A⟶B), but without any immediate reversion (B⟶A). A full description of the definitions used here for homoplasy and/or stepwise evolution are available as Supplementary Text 1 and Supplementary Figure 1. A full description of the basic steps used in our algorithm, including a schematic representation, is available in the Supplementary Text 2, Supplementary Figure 2, and Supplementary Figure 3. Associated code is publicly available at https://github.com/nataliamv/SARS-CoV-2-ARTs-Classification.

### 4. Estimating dN/dS

Derived from the global alignment and tree, we estimated dN/dS (ω, the ratio between the non-synonymous substitution rate per non-synonymous site and the synonymous substitution rate per synonymous site) using site, branch and branch-site models: Mixed Effects Model of Evolution (MEME), Fixed Effects Likelihood (FEL), and the fixed effects site-level model (Contrast-FEL) (Kosakovsky Pond and Frost 2005; Murrell et al. 2012; Kosakovsky Pond et al. 2021). For this, the alignment was partitioned into 14 putatively non-recombinant regions using the Genetic Algorithm for Recombination Detection (GARD) (Kosakovsky Pond, Posada, et al. 2006), with all subsequent analyses conducted on the partitioned data. As dN/dS models use the GTR component for the nucleotide evolutionary rate, biased mutation rates are handled. Further, to mitigate the inflation in dN/dS estimates that results from unresolved and/or maladaptive evolution, testing for selection was again restricted to internal nodes/branches of the phylogeny (Kosakovsky Pond, Frost, et al. 2006). Genome-wide comparison of dN/dS estimates across viral genome regions was performed using the Branch-Site Unrestricted Statistical Test for Episodic Diversification method (BUSTED) (Murrell et al. 2015). Finally, the impact of changing biochemical properties at selected sites was further assessed under the Property Informed Models of Evolution method (PRIME) (HyPhy 2013). Our results were further compared to the selection analysis available for independently sampled SARS-CoV-2 genome data available as of December 2022 (https://observablehq.com/@spond/evolutionary-annotation-of-sars-cov-2-covid-19-genomes-enab) (Kosakovsky Pond).

### 5. Mapping mutations onto betacoronavirus protein structures

To locate the non-synonymous mutations identified on viral protein regions of known function, corresponding residues were mapped to available structural data using PyMOL v 2.4.0 (https://pymol.org/2/) (Supplementary Table 3, see Data Availability section). Mutations were analysed in the context of their relative proximity to previously reported functional regions, and to each other. N-linked glycosylation sites in S protein sequences were identified by searching for the N-[not P]-[S or T] consensus sequence (Watanabe et al. 2019). None of the mutations identified in this study resulted in generation or deletion of N-linked glycosylation sequons. In parallel, the identified conserved and variable sites as well as the 30 mutations evidencing homoplasy and/or stepwise evolution across virus species were mapped onto published protein structures available for the S proteins of the four human-infecting betacoronaviruses studied here (Figure 5, Supplementary Figure 7). Finally, to compare dN/dS distributions between specific domains of the S protein within and across virus species, sites inferred to be under positive or negative selection (PSS, NSS) were mapped onto S protein structures (Supplementary Data 2).

### 6. Validation through resampling and by comparing mutational distributions

To validate our initial observations derived from virus genomes sampled up to May 2021, we sought to determine if the 30 mutations that had been identified initially were also present in the expanded embecov- and sarbecovirus diversity sampled up to July 25^th^ 2022 (corresponding to the final sampling date of this study). Virus diversity now included genome sequences derived from more recently collected human isolates (only made publicly available after our initial sampling), and from other closely related embeco- and sarbecoviruses from non-human hosts. The expanded alignment comprises 1455 sequences (∼700 embecovirus + SARS-CoV and ∼700 SARS-CoV-2), resulting in 27503 columns that were re-aligned under a progressive profile-to-profile approach based on taxonomic relatedness to be further used to estimate an expanded ‘Maximum Likelihood’ tree (following Methods section 2). To additionally explore if the mutations identified were also present in a larger dataset representing an expanded SARS-CoV-2 diversity (sampled up to July 25^th^ 2022), a set of 1400 SARS-CoV-2 genomes denoting ‘evolutionary successful’ virus lineages (Supplementary Table 1) was examined independently (Supplementary Text 3, Supplementary Figure 5). Both datasets were analysed following the steps described in Methods Section 2 and 3, specifically searching for the mutations listed in Table 1. Virus taxa included in both re-sampled datasets are listed in Supplementary Data 5. A full description of the subsampling methodological approach used is available as Supplementary Text 3, and Supplementary Figure 4.

We further sought to explore if the proportion of mutational patterns we classified as putatively denoting homoplasy and/or stepwise evolution were more likely to arise from an evolutionary scenario mostly driven by selection, compared to ‘random’ mutational patterns derived from evolutionary scenarios generally driven by genetic drift. For this purpose, the expanded alignment was translated to amino acid sequences and used to simulate three alignments with ‘AliSim’ (http://www.iqtree.org/doc/AliSim) under the ‘mimick real alignment’ function (mimicking a ‘real’ evolutionary process based on amino acid evolution under a LG model, and applied to the inputted original tree). To compare the corresponding proportion of sites scored under homoplasy and/or stepwise evolution, each dataset (the expanded and three simulated alignments) was analysed following the steps described in Methods section 3. The classification of mutation patterns within expanded and simulated datasets also serves the purpose of validating our algorithm, originally developed for analysing the global dataset (including merely OC43, HKU1, SARS-CoV-1 and SARS-CoV-2). Associated results and a brief discussion are available as Supplementary Text 5.

### 7. Reconstruction of amino acid evolution for selected sites

For those mutations displaying cumulative evidence of being potentially informative on adaptive convergence (sites 18121, 21623 and 23948) (Table 1), we used the expanded dataset to infer ancestral states under a Bayesian framework. For this, we first estimated an MCC (maximum clade credibility) tree from the resampled codon alignment using a SRD06 substitution model (Shapiro et al. 2006) and a strict molecular clock. For each site of interest, coded amino acid traits were mapped onto the nodes of the MCC tree by performing reconstructions of ancestral states under an asymmetric discrete trait evolution model (DTA) in BEAST v1.8.4 (Lemey et al. 2009; Suchard et al. 2018). The DTA model was run using a Bayesian Skygrid tree prior for 100×10^6^ generations and sampled every 10,000 states until all DTA-relevant parameters reached an ESS >200. The resulting trees were used as an input for Figure 3.

## ACKNOWLEDGEMENTS

MEZ was supported by Leverhulme Trust ECR Fellowship (ECF-2019-542). RJGH is supported by the European Molecular Biology Organisation (ALTF 869-2019). RJGH and TAB are supported by the Medical Research Council (MR/S007555/1). The Wellcome Centre for Human Genetics is supported by Wellcome Trust grant 203141/Z/16/Z. SKP is supported in part by the the NIH (grants AI134384, AI140970, GM110749) and the NSF (grant 2027196). JT and RPDI were supported by European Union’s Horizon 2020 project MOOD (874850). LvD is supported by a UCL Excellence Fellowship. HGCS is supported by funding through the “Vigilancia Genómica del Virus SARS-CoV-2 en México” grant from the National Council for Science and Technology-México (CONACyT). Author contributions: MEZ and OGP designed research. MEZ and RJGH performed research. MEZ, RJGH, BG, SKP, LvD, JT, RPDI and HGCS analysed data. NM developed the code for implementing the computational pipeline. OGP and TAB supervised data analysis. MEZ and RJGH wrote the manuscript, with comments from all authors. We thank Dr. Louis Du Plessis for his help with the phylogenomic analysis and Dr. Nicola De Maio for constructive comments on our analytical approach.

## COMPETING INTERESTS

The authors declare no competing interests.

## DATA AVAILABILITY STATEMENT

Taxa IDs and accession numbers for virus sequences used in this study are provided in the Supplementary Data 4 and 5 files. All SARS-CoV-2 genome sequences and associated metadata used in this study are published in GISAID’s EpiCoV database under the EPI SET GISAID Identifier: EPI_SET_230131zy. To view the contributors of each individual sequence with details such as accession number, virus name, collection date, originating and submitting lab, as well as the list of all authors, visit <u>10.55876/gis8.230131zy.</u> PBD files used are listed as follows: S protein (HKU1 PDB:5I08, OC43 PDB:6OHW, SARS-CoV-1 PDB:6ACC and SARS-CoV-2 PDB:6VXX, 6ZGI). Orf1a (SARS-CoV-1 nsp3 PDB:2W2G). Orf1b (SARS-CoV-2 nsp13 PDB:6XEZ, SARS-CoV-1 nsp14 PDB:5C8S and SARS-CoV-2 nsp15 PDB:6WLC). Full code for our algorithm is available as open source: https://github.com/nataliamv/SARS-CoV-2-ARTs-Classification. An interactive notebook with our full selection analysis results is available at https://observablehq.com/@spond/beta-cov-analysis.

## Notes

https://github.com/nataliamv/SARS-CoV-2-ARTs-Classification

https://observablehq.com/@spond/beta-cov-analysis

### Competing Interest Statement

The authors have declared no competing interest.

### Summary of Updates

Since our manuscript has been available as a preprint (May 2021), we have made substantial changes to improve its quality and fulfil the high-quality standards set by the journal. This includes validation of our initial observations pursued through an expanded resampling of virus diversity (available up to July 25th 2022, corresponding to the final sampling date of this study). In this light, some of the mutations we initially detected are now known to display dominant trends in SARS-CoV-2. Moreover, based on comparing mutational distributions between experimental and simulated data, we confirm that the proportion of amino acid trait evolution patterns observed within the experimental data were more likely to arise from evolutionary scenarios driven mostly by selection, compared to random mutational patterns driven by genetic drift (represented by the simulated data).

