## Supplementary Information for "Identification of evolutionary trajectories shared across human betacoronaviruses"

### SUPPLEMENTARY INFORMATION
### "Identification of evolutionary trajectories shared across human betacoronaviruses"

#### INTRODUCTION

**Supplementary Text 1.** Definition of homoplasy and stepwise evolution

**Supplementary Figure 1.** Patterns of amino acid evolution potentially informative on adaptation

#### METHODS AND RESULTS

**Supplementary Text 2.** Methods: description of the algorithm for classifying genetic variation patterns as putative homoplasy and stepwise evolution

**Supplementary Figure 2.** Basic steps followed in the algorithm

**Supplementary Figure 3.** Confusion matrix comparing the classification of ARTs according to different categories (homoplasy/stepwise evolution/others) under visual inspection or under our algorithm

**Supplementary Text 3.** Methods: validation through an expanded re-sampling of the known betacoronavirus diversity

**Supplementary Table 1.** Variant of Concern (VOC), Variant under monitoring (VUM), or Variant of Interest (VOI) designation reported by the WHO up to July 25th 2022

**Supplementary Figure 4.** Distribution of SARS-CoV-2 genome sequences before and after subsampling

**Supplementary Figure 5**. ML comprising the re-sampled 1400 SARS-CoV-2 sequences

#### RESULTS

**Supplementary Text 4.** Results: structural and functional contextualization of other mutations identified to denote homoplasy and/or stepwise evolution

**Supplementary Text 5.** Results: comparison of the proportion of sites scored under homoplasy and/or stepwise for the expanded and simulated datasets

**Supplementary Figure 6.** Position of a variable residue (genome site 23948) within the region directly preceding the S2' cleavage site.

**Supplementary Figure 7.** Distribution of positively (PSS) and negatively (NSS) selected sites within the SARS-CoV-2 Spike protein structure

**Supplementary Table 2.** PSS identified across all four virus species studied

**Supplementary Table 3.** Location of other sites displaying evolutionary patterns putatively denoting homoplasy/stepwise evolution on protein regions of known function

#### SUPPLEMENTARY DATA

- **Supplementary Data 1.** Maximum likelihood phylogenies obtained from the individual and global alignments estimated using RAxML (Supplementary_Data_1_ML_treefiles.pdf)
- **Supplementary Data 2.** Distribution of conserved sites across S and site-specific dN/dS values (Supplementary_Data_2_Conserved_Sites.xlsx)
- **Supplementary Data 3.** Reconstruction of ancestral states on the base RAxML tree under ML approach (RAS-ML) (Supplementary_Data_3_RASML_Orf1ab_S.pdf)
- **Supplementary Data 4.** Taxa IDs and accession numbers for sequences used, and GISAID acknowledgements for the COG dataset (Supplementary_Data_4_Taxa_ID.xlsx)
- **Supplementary Data 5.** Taxa IDs and accession numbers for the 'expanded alignment' and the 're-sampled SARS-CoV-2 genomes' datasets

**Supplementary Text 1. Definition of homoplasy and stepwise evolution**

Differential selective pressures exerted by the host immune responses impact the phylogenetic patterns of viral populations (Grenfell et al. 2004), an observation of particular importance when exploring the effects of positive selection applied to amino acid trait evolution. Two patterns that may arise after positive selection are *homoplasy* and *stepwise evolution*. Homoplasy (including both parallel and convergent evolution) is defined as the appearance of comparable traits (i.e., mutations) in lineages that do not share direct common ancestry, and has been described as a well-documented hallmark of adaptation for different RNA viruses (Gutierrez et al. 2019). For example, the parallel loss of the hemagglutinin-esterase (HE) protein lectin function in endemic betacoronaviruses (HKU1 and OC43) likely reflects convergent adaptation to the human hosts (Bakkers et al. 2017). Moreover, recurring parallel mutations have been linked to the emergence of highly-pathogenic viral genotypes/phenotypes (Stern et al. 2017; Escalera-Zamudio et al. 2020). Conversely, stepwise evolution is represented by sequential mutations resulting in novel genotypes that may denote 'evolutionary success', shifting towards new local optimums on a changing fitness landscape (Dolan et al. 2018). The accumulation of sequential mutations over time may reflect the effects of selective pressure exerted by long-term host immune responses, denoting evolutionary processes such as antigenic drift (Boni et al. 2006; Starr et al. 2021), and the adaptation to novel ecological niches, such as those represented by host switching events (Menachery et al. 2017). Following from Dollo's parsimony principle (stating that once a complex trait is lost during evolution, it is unlikely to be regained) (Farris 1977; Delport et al. 2008), stepwise evolution can be defined as consecutive mutational changes between different states (A→B, then from B →C), without any immediate reversions (B→A) (Supplementary Figure 1).

**Supplementary Figure 1. Patterns of amino acid evolution potentially informative on adaptation**

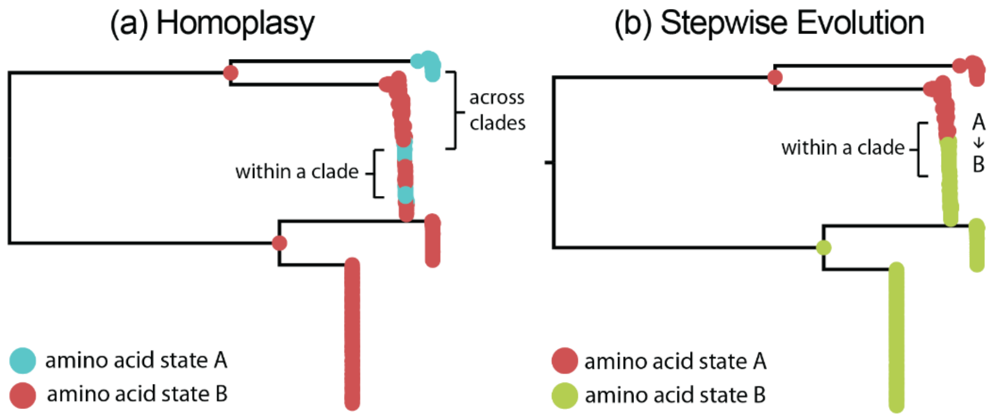


Mutational patterns that may be informative on adaptation include homoplasy and stepwise evolution. (a) Homoplasy can occur within a single clade and/or across clades, in which the same amino acid change is observed in at least one internal node of a given clade, and in another internal node of the same or a different clade. Clades with shared amino acid states must not share direct common ancestry. (b) Stepwise evolution can occur only within a single clade, and is defined as sequential mutations observed to occur at the same site across time, displaying transitional changes between different states (A→B), but without immediate reversions (B→A). Homoplasy and stepwise evolution are not mutually exclusive, and may co-occur.

**Supplementary Text 2. Methods: description of the algorithm for classifying genetic variation patterns as putative homoplasy and stepwise evolution**

The algorithm takes as input the Ancestral Reconstruction Trees (ARTs) in a NEXUS format derived from the amino acid evolution analysis (RASML) performed on a given ML tree (Sagulenko et al. 2018) , in which inferred ancestral amino acid states derived from the codon alignment are coded as discrete traits and mapped onto the nodes of the tree. RASML analysis of the global alignment resulted in 6,681 ARTs, further inputted into our algorithm developed to classify amino acid change patterns putatively denoting homoplasy and stepwise evolution compared to 'other' and/or 'conserved' mutational patterns (see Supplementary Text 1 and Figure 1). The algorithm consists of two parts: first, for each ART, a pre-processing stage eliminates redundancy without losing information. Second, amino acid evolution patterns are classified according to 'the patterns of interest' (in this case, homoplasy and/or stepwise evolution). Homoplasy can occur at an inter- or intra-clade level, defined as any given amino acid change occurring in at least one internal node of a given clade that must also be present in at least another internal node of the same or another clade. Nodes with a shared amino acid state must not directly descend from a common ancestor node. Stepwise evolution can occur only within the same clade, defined here as any sites displaying directional mutations involving at least two states (A→B), but without any immediate reversions to the ancestral state (B→A). The script was built using the Phylo module in the BioPython package (Cock et al. 2009) to parse, modify and export reduced ARTs to the NetworkX package (A et al. 2008). NetworkX was used in the main part of the algorithm to identify different patterns of amino acid evolution, as it allows flexibility in manipulating network-like objects. A combination of NetworkX and Matplotlib (Hunter 2007) was used to visualise the resulting ARTs. The open-source code is fully available at https://github.com/nataliamv/SARS-CoV-2-ARTs-Classification. The script consists of the following basic steps (see Supplementary Figure 2):

**PRE-PROCESSING**:

a) Preliminary information is collected from the topology of the inputted ML tree, including tip names (*i.e.* Taxa IDs). Since all ARTs derive from the same 'base' ML tree, they are initially equivalent. However, although all ARTs share the same structure, amino acid states observed at different nodes may vary. We first defined the distance between two nodes as the number of branches between them, so that the distance between a descending node and its immediate ancestor node is always equal to one. *K* is defined as the distance from any node to the root of the tree. The global tree of human-infecting betacoronaviruses has four different branches leading to four main clades (HKU1, OC43, SARS-CoV-1 and SARS-CoV-2), with a distance from the root node to the common ancestor of each one being $K = 2$.

b) Each ART is then read into memory and reduced in the following way:
   - **a.** Tips are collapsed to their most recent ancestral node. Only internal nodes are kept, and a name and weight attribute of 1 is added to each node, in order to preserve information.
   - **b.** Each ART is once more collapsed according to the amino acid state present at each node as follows: a given node 'B' is merged with its immediate ancestor 'A', if all descending nodes share the same amino acid state (or if the state of the node is 'X'). Then the weight value of node 'B' is added to the value of ancestor node 'A', with the amino acid state observed for a 'A' being preserved. In this way, weight values of newly merged nodes represent the number of consecutive merging events of equal states. The process is carried out in an iterative way starting at the terminal nodes and moving towards the root, stopping at the nodes at a K+1 distance from the root node. In this way, the general divergence pattern observed for the base ML tree is preserved.
   - **c.** For each ART, the weight attribute (w) of each terminal node is checked. Whenever w<=5, the ART is then flagged as containing nodes representing 'Low Frequency Variants', that are to be removed from the given ART.

c) Each reduced ART is exported to NetworkX and saved using the DiGraph class (graphs with directed edges). Given the binary nature of the basic pattern of any phylogenetic tree (*i.e.* each node has two descendants), saving ARTs as DiGraphs allows us to access the ancestry-descendance information of the tree in order to effectively identify ancestors. This is given the structural properties of tree-like digraphs, such as the absence of cycles and the existence of a unique path between each pair of nodes.

**CLASSIFICATION:**

d) Once pre-processing is completed, we sought to identify and filter out all reduced ART (r-ART) with patterns representing conserved states (*i.e.* all nodes share the same state). To achieve this, the amino acid states present at each node of the r-ART were added to a 'set' (S). A set is defined as an unordered collection with no duplicate elements. In this sense, if the number of elements in *S* = 1, the amino acid states observed for all nodes within the r-ART is the same. In this case the r-ART is classified as conserved, and the execution of the algorithm stops. If the r-ART is not classified as 'conserve', it is then further tested for 'Homoplasy' and/or 'Stepwise' evolution as follows:

a. **Homoplasy**:

Homoplasy is identified at an inter-clade level (across clades) and intra- (within a single clade):

- Homoplasy at an inter-clade level. States shared across clades are saved to two different sets. In this case, we use S1 for clades 1 and 2 (*i.e.,* HKU1 and OC43), and S2 for clades 3 and 4 (*i.e.*, SARS-CoV-1 and SARS-CoV-2). We then construct a new set corresponding to the intersection of S1 and S2, which includes all states shared by S1 and S2:

    1. If the intersection of clades S1 and S2 is empty (*i.e.* no nodes within different clades share the same state), then inter-clade homoplasy is ruled out.
    2. If the intersection of clades S1 and S2 is not empty (*i.e.* at least a pair of nodes within different clades share the same state), two additional scenarios follow:
        I. If the most recent ancestral nodes of each clade share the same state, then inter-clade homoplasy is ruled out.
        II. If the most recent ancestral nodes of each clade do not share the same state and both sets have more than one state, then inter-clade homoplasy is scored.

- Homoplasy at an intra-clade level. For each clade, all terminal-to-root node paths are analysed. Consecutive nodes with equal states are merged, and if the resulting path has two equal states appearing more than once, then homoplasy at intra-clade level is scored.

b. **Stepwise evolution:**

For classifying patterns under stepwise evolution, all terminal-to-root node paths are analysed. Equal consecutive states are merged, and if the resulting path has two or more different consecutive states, then the pattern is classified as stepwise evolution.

e) Each ART is assigned a vector of length 4, with 1 or 0 recording presence/absence of patterns denoting conserved states, inter-clade homoplasy, intra-clade homoplasy or stepwise evolution. The vectors are stored and exported as a csv file.

Comparing the classification of ARTs according to different categories ('Homoplasy/Stepwise' versus 'Conserved/Others') under visual inspection or by using the algorithm, revealed that the algorithm is more sensitive identifying ARTs under 'Homoplasy/Stepwise' than the visual inspection (Supplementary Figure 3). The number ARTs classified under 'Homoplasy/Stepwise' or' Others' under both methods is consistent.

**Supplementary Figure 2. Basic steps followed in the algorithm**

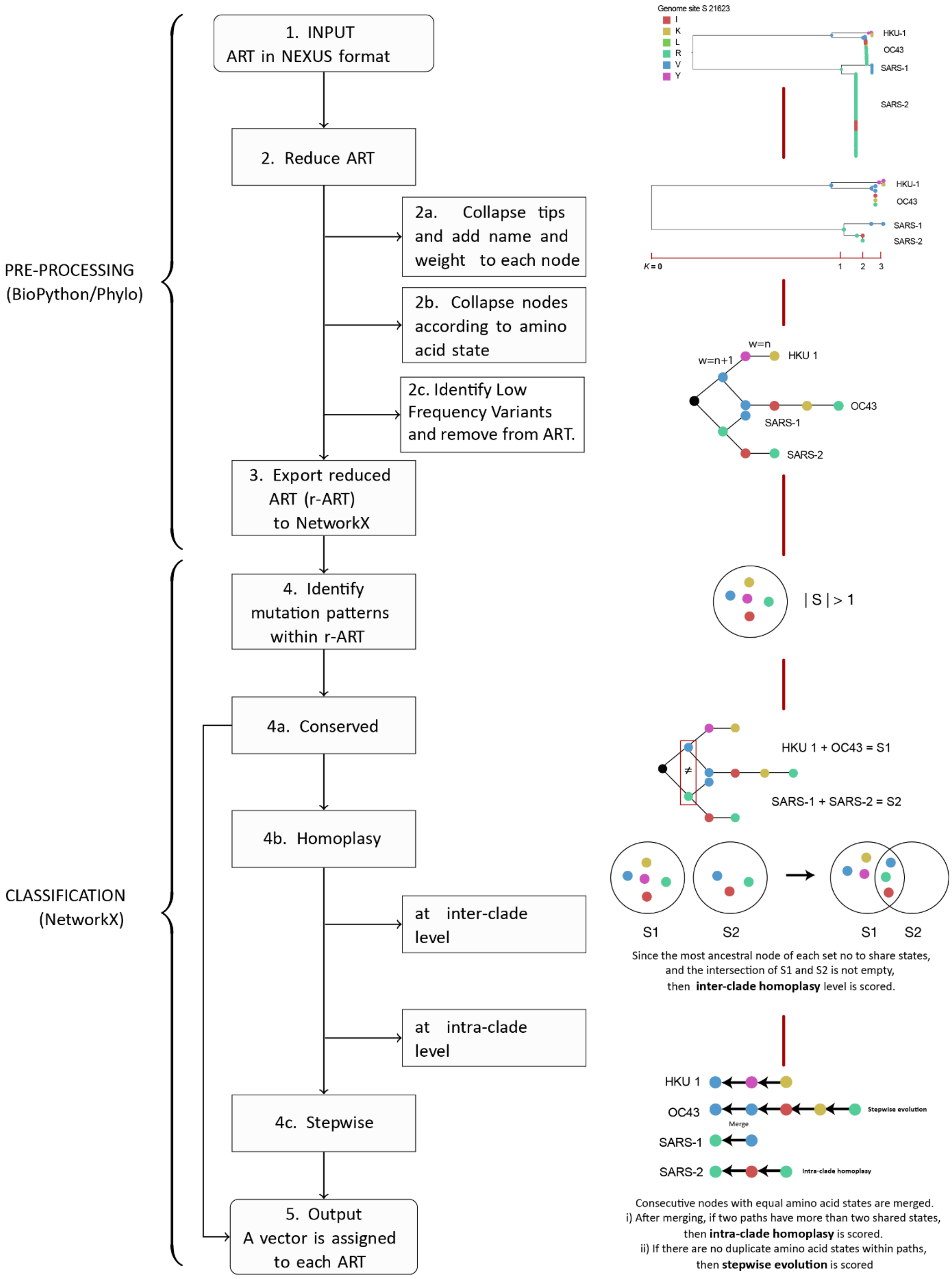


ART pre-processing is carried out in order to reduce complexity, but without losing information (represented by names and weight attributes for each nodes). The diagram represents the sequential steps implemented within our algorithm used for classifying genetic variation patterns within ARTs according to the different evolutionary patterns ('Homoplasy' and/or 'Stepwise' evolution).

**Supplementary Figure 3. Confusion matrix comparing the classification of ARTs according to different categories (homoplasy/stepwise evolution/others) under visual inspection or under our algorithm**

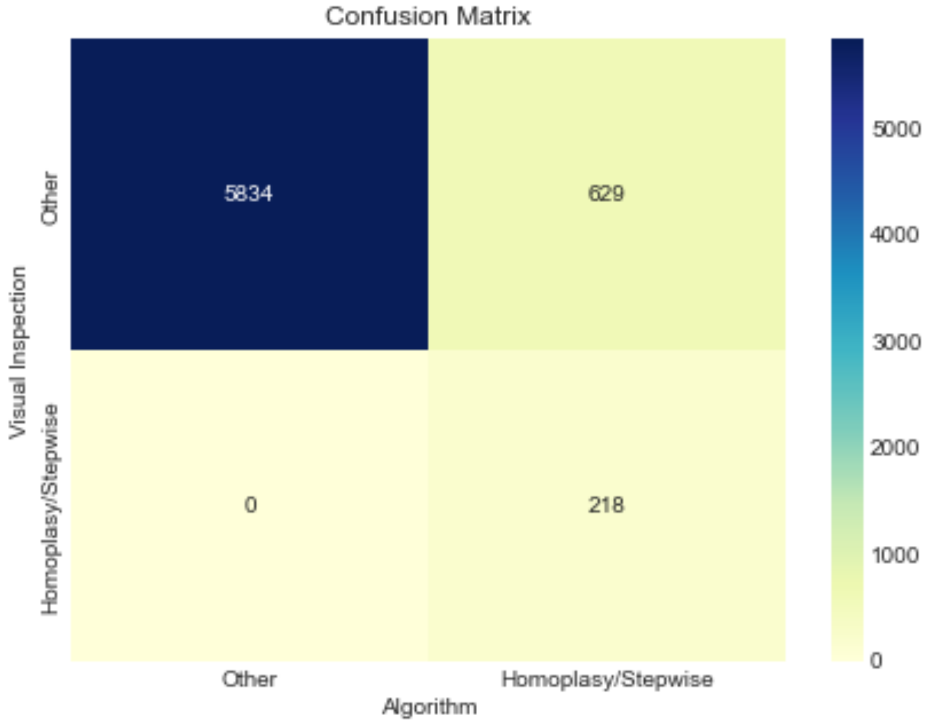


Each panel represents the number of coincidences/disagreements for the number of ARTs classified either visually or using the algorithm. The top left panel indicates number of coincidences for ARTs classified as ‘Others’, whilst lower right panel indicates number of coincidences for ARTs classified as ‘Homoplasy/Stepwise’. The colour scale indicates the number of ARTs classified under each category.

**Supplementary Text 3. Methods: validation through an expanded re-sampling of the known betacoronavirus diversity**

To generate the expanded dataset, approximately 450 embecoviruses (including HKU1, OC43, and related viruses from animal hosts), 80 sarbecoviruses (including SARS-CoV-1, and related viruses from animal hosts; excluding SARS-CoV-2), and 190 MERS-CoV (and MERS-like) complete viral genomes were retrieved from the Virus Pathogen Resource (ViPR-NCBI 2021). These sequences represent all publicly available virus genomes sampled in different countries and from different host species across time, available up to July 25th 2022 (corresponding to the final sampling date of this study). Genomes were filtered according to the quality criteria stated in Methods section 1 (main text), whilst potential recombinant sequences were identified and removed using ClonalFrameML (Didelot and Wilson 2015). In total, less than 10 sequences were removed for the OC43 viruses (C, E, F and G genotypes) (Oong et al. 2017). Non-recombinant fragments were verified using GARD (Kosakovsky Pond et al. 2006). In total, ~700 genome sequences from all virus species were retained. MERS-CoV sequences were included for tree rooting purposes, but excluded from all subsequent analyses.

To generate the re-sampled SARS-CoV-2 dataset, the sequence sub-sampling approach we used was based on the following steps:

- Selecting only SARS-CoV-2 diversity representing ‘evolutionary success’: virus lineages and sub-lineages designated as Variant of Concern (VOC), Variant under monitoring (VUM), or Variant of Interest (VOI) (WHO 2022)(add REF) known until the final sampling date of this study (Supplementary Table 1).
- A uniform temporal sampling across epidemiological weeks (Inward, Parag, et al. 2022; Inward, Jackson, et al. 2022), but proportional to the global representation of each virus lineage selected (Supplementary Figure 2),
- A phylogenetically-informed subsampling used to further reduce genetic overrepresentation with a minimal loss of diversity, thus preserving the overall phylogenetic clustering patterns reported for SARS-CoV-2 (Menardo et al. 2018; Escalera-Zamudio et al. 2020; Hill et al. 2021). The earliest viral genomes dating to before any variant designation were retained and labelled as ‘unclassified’, kept for tree rooting purposes.

Virus genomes available in the GSAID platform up to July 25th 2022 (GISAID 2021) were downloaded and further filtered using the quality control pipeline available in the ‘Nextclade’ platform (https://clades.nextstrain.org/). Sequences were assigned to their corresponding PANGO lineage and variant designation using the Pangolin v.4.1.2 software tool (https://pangolin.cog-uk.io/), to be further subsampled.

Following the steps abovementioned, a set of approximately 10,000 subsampled genome sequences was obtained and aligned by mapping to the reference genome (Wuhan-Hu-1, GenBank: MN908947.3) using Minimap2 (Li 2018). Subsequently, the main viral ORFs (Orf1ab and Orf S) were extracted to generate a reduced-length alignment of approximately 25,000 bases long, used to estimate a local Maximum Likelihood (ML) tree using IQ-TREE (command line: iqtree -s -m GTR+I+G -alrt 1000) (Minh et al. 2020). The resulting tree was then further reduced using the phylogenetically informed approach Treemmer v0.3 (Menardo et al. 2018), resulting in a ML tree comprising approximately 6500 SARS-CoV-2 sequences. However, the number of SARS-CoV-2 sequences had to be further reduced to be equated to the total number of embecovirus + sarbecovirus sequences available in the expanded dataset, as an unbalanced number of taxa could distort the tree topology (Holmes and Rambaut 2004).

Therefore, following a secondary phylogenetically-informed subsampling iteration, the 6,500 SARS-CoV-2 genomes dataset was again reduced into two secondary datasets: one of 700 and one of 1,400 sequences. The set of 700 sequences was added to the expanded alignment, resulting in a total of 1,455 genome sequences comprising the four virus species studied. The dataset representing the re-sampled 1,400 SARS-CoV-2 sequences was inputted for independent analyses. In this context, the analysis of re-sampled virus diversity can further help detect mutations erroneously scored as ‘shared’ as a result of alignment uncertainty for hypervariable regions (common across highly divergent taxa). Moreover, adding viruses from non-human hosts may help mitigate the potential effects of long branches, facilitating the identification of homologous mutations within deeper nodes of the tree (Holmes and Rambaut 2004).

**Supplementary Table 1. Variant of Concern (VOC), Variant under monitoring (VUM), or Variant of Interest (VOI) designation reported by the WHO up to July 25th 2022**

| **VOC** | |
|---|---|
| Omicron | B.1.1.529 |
| Alpha | B.1.1.7 |
| Beta | B.1.351 |
| Gamma | P.1 |
| Delta | B.1.617.2 |
| **VOI** | |
| Epsilon | B.1.427 and B.1.429 |
| Zeta | P.2 |
| Eta | B.1.525 |
| Theta | P.3 |
| Iota | B.1.526 |
| Kappa | B.1.617.1 |
| Lambda | C.37 |
| Mu | B.1.621 |
| **VOM** | |
| AV.1 | B.1.619 |
| AT.1 | B.1.620 |
| R.1 | B.1.630 |
| B.1.466.2 | B.1.1.318 |
| B.1.1.519 | C.1.2 |
| C.36.3 | B.1.640 |
| B.1.214.2 | XD |
| B.1.1.523 | |

**Supplementary Figure 4. Distribution of SARS-CoV-2 genome sequences before and after subsampling**

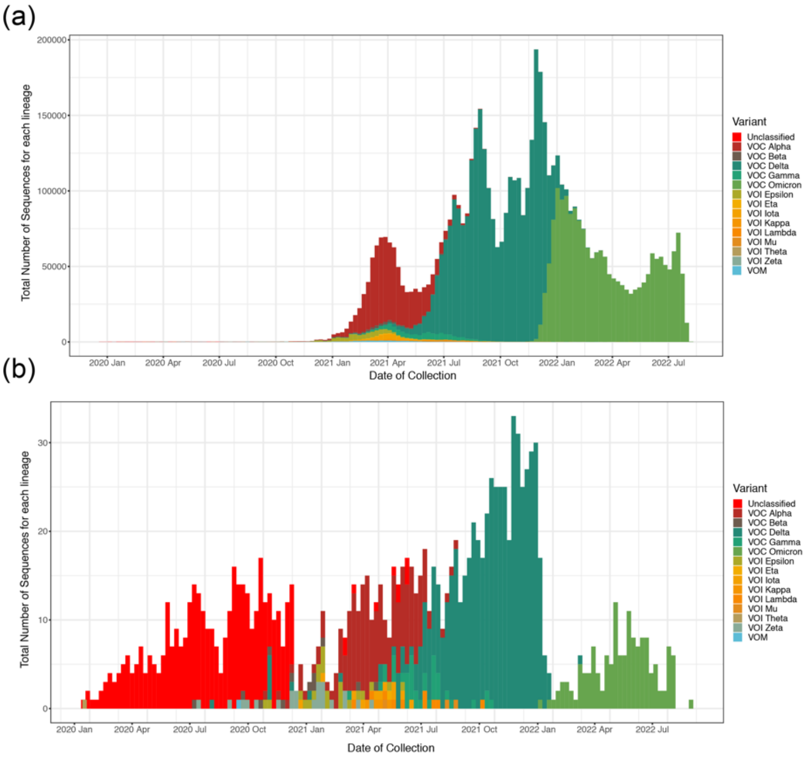


Distribution plot (a) showing VOC/VOI/VUM sequences sampled across time (from April 2020 up to July 2022), whilst plot (b) shows the proportional number of the different VOCs/VOIs/VUMs subsampled across epidemiological weeks (resulting in 62 sequences per week). Sampled genomes are coloured according to their VOC/VOI/VUM designation.

**Supplementary Figure 5. ML comprising the re-sampled 1400 SARS-CoV-2 sequences**

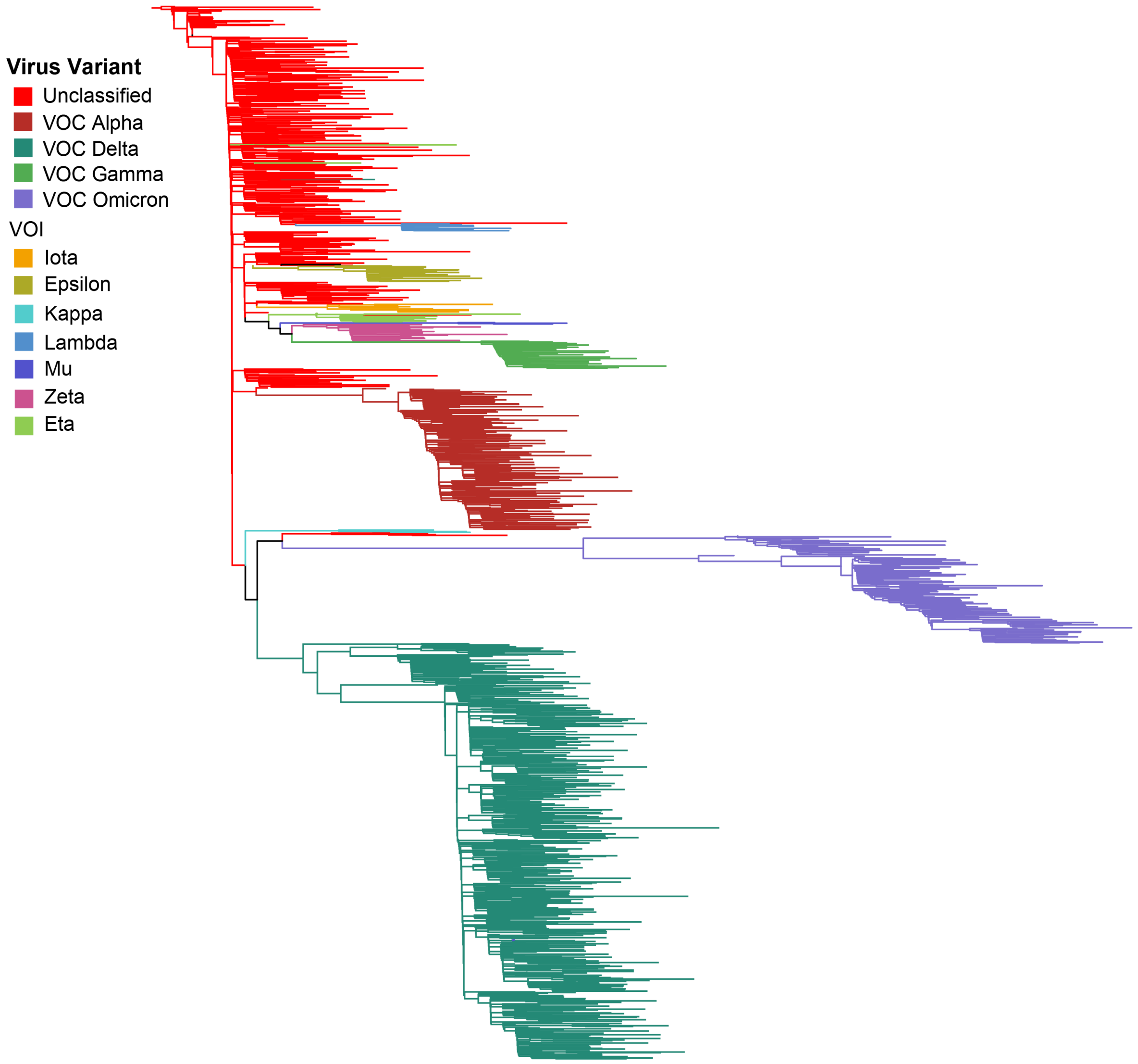


Maximum Likelihood tree estimated from the Orf1ab+S alignment derived from the re-sampled 1,400 SARS-CoV-2 sequences (used for further validation analyses, see Supplementary Text 3). The tree summarizes the phylogenetic relationship observed for current SARS-CoV-2 diversity (sampled up to July 25$^{th}$ 2022), highlighting virus lineages representing 'evolutionary success' [Variant of Concern (VOC), Variant under monitoring (VUM), or Variant of Interest (VOI)].

**Supplementary Text 4. Results: structural and functional contextualization of mutations identified to denote homoplasy and/or stepwise evolution**

To gain insight into the functional relevance of sites/mutations identified as displaying evidence for homoplasy and/or stepwise evolution listed in Table 1 (main text), we searched the literature for functional regions within residue proximity. Our findings are described below, excluding sites 18121, 21623, 21635 and 23948 that are thoroughly discussed in the Results section of the main text. We would like to stress that additional functional regions of the betacoronavirus proteome may be identified in the future, and thus our current analysis may therefore miss out other potentially relevant mutations. In addition to sites 18121, 21623, 21635 and 23948, we find 4 more sites/mutations that are proximal to regions of known protein function (20344 in nsp15, and 24614, 24620 and 24632 in S2), while also displaying evidence of homoplasy/stepwise evolution and positive selection. These are summarized in Supplementary Table 3 and are discussed below:

- Site 20344 also in Orf1ab corresponds to a non-conservative mutation 'H' to 'Y' that is observed to be homoplasic in some HKU1 and sarbecoviruses (Supplementary Data 3, Table1). This site corresponds to residue 243 in nsp15 (according to the SARS-CoV-2 protein numbering) (Supplementary Figure 5), located within the NendoU catalytic domain of the endoRNAse. This domain specifically targets and degrades viral mRNA polyuridine sequences to prevent host innate immunity signalling (Hackbart et al. 2020). However, this mutation is located distally (~12 Å) to the nucleotide binding pocket of the active site, and the potential impact upon endoRNAse activity is unknown.
- Sites 24614, 24620 and 24632 in S2 correspond to the conservative mutation 'V' to 'I' at residue 1018, 'F' to 'A' at residue 1020, and the non-conservative mutation 'L' to 'R' at residue 1024 in S2 (according to the SARS-CoV-2 protein numbering), observed to be homoplasic for some HKU1 and sarbecoviruses. These three residues are within close proximity of each other (the distance between individual sites is less than 10 Å), positioned within the central helix of S2 (Figure 5, main text), a region which conformational rearrangements facilitate membrane fusion (Kirchdoerfer et al. 2016; Pallesen et al. 2017; Kirchdoerfer et al. 2018) Again, given their close proximity (and possible epistatic interactions), it is likely that these amino acid changes may collectively alter the fusogenic functionality of the S2 subunit, as has been observed for other changes located within the central helix domain of coronaviruses (Hulswit et al. 2016).

The remaining sites identified in this study to putatively denote homoplasy and/or stepwise evolution (see Table 1) were not found to be near currently recognised functional protein regions of betacoronaviruses. To a certain extent, this may be due to incomplete appreciation of betacoronavirus protein function and structure. The structural contextualization for all 26 sites/mutations listed in Table 1 that are not discussed in the main text, is described below. To maintain a more complete overview, the four sites described in the above Supplementary Text (20344, 24614, 24620, 24632) are included here as well.

**Orf1ab**

- Site 2557 maps to a region on nsp2 distal to the highly-conserved zinc ion-binding site that is suggested to be involved in RNA binding (Gupta et al. 2021).
- Site 7478 resides in nsp3 (residue 1578), recently shown by cryo-electron tomography to form a molecular pore that spans the double membrane of coronavirus replication organelles (Wolff et al. 2020) . This site appears to map to a region within one of the transmembrane domains of the molecular pore.
- Genomic sites 16189 and 17809 map onto nsp12 (RNA-dependent RNA polymerase) and nsp13 (helicase). Together form the core of the coronaviral replication transcription complex (Chen et al. 2020).
- Sites 18334, 18442 and 19048 map to different domains of the nsp14 endoribonuclease (Ma et al. 2015). Sites 18334 and 18442 are found in close proximity of one another (~7 Å) in the ExoN domain, but are located further away from site 18121 described in the main text (>15 Å). Site 19048 is located in the methyltransferase domain.
- Sites 20344 and 20554 map to the NendoU catalytic domain of the nsp15 endoRNAse (Kim et al. 2021). They are part of the same face of this domain on the side of the uridine binding pocket, but not in close proximity of one another (~25 Å) or the uridine binding pocket itself (>12 Å).
- Site 21400 maps to the surface of the nsp16 2'-*O* methyltransferase (Rosas-Lemus et al. 2020).

**Orf S**

For the 16 remaining sites identified within S, 9 ($S1^A$: 21614, 21800, 21863, 21920, 21926, 22004 and 22124; $S1^B$: 22553 and 23048) mapped onto S1 and 7 onto the S2 subunit (24614, 24620, 24632, 24863, 25037, 25166 and 25247) (Walls et al. 2020). At the moment, there are no near atomic level structures available for mutations corresponding to genomic sites 25037, 25166 and 25247 (S), and thus these cannot be further analysed.

**Supplementary Text 5. Results: comparison of the proportion of sites scored under homoplasy and/or stepwise for the expanded and simulated datasets**

An overview of the simulated data revealed that the three replicates were almost identical to each other, enabling us to compare average values with the expanded data. General alignment statistics before masking conserved sites revealed that the simulated alignments displayed a mean pairwise identity of approximately 40% compared to a 60% observed for the expanded alignment. After retaining variable sites only (see Methods section 3, main text), the simulated alignments displayed a mean pairwise identity of 40% compared to 50% observed for the expanded alignment. These observations suggest that the expanded alignment displays fewer variable sites, possibly linked to the effects of purifying selection reducing the number of non-synonymous mutations observed on the experimental (real) data.

When analysing simulated data under our algorithm for the classifying patterns of genetic variation to detect putative homoplasy and/or stepwise evolution (see Methods section 3 in main text, as well as Supplementary Text 2, Supplementary Figures 2 and 3), we noted that the number of sites scored under ‘homoplasy' within the simulated data was generally lower compared to the expanded data. For example, only 1% of all sites were scored under both intra- and inter-clade homoplasy for the simulated data, whilst an estimate of 3% was observed for the expanded data. Similarly, an estimate of 3% versus 7% was observed for all sites scored as intra-clade homoplasy. This suggest that despite the simulated data displaying more variable sites, a reduced proportion of mutational patterns follow putative homoplasy, further evidencing that the sites we detect within the experimental (real) data are more likely to have resulted from an evolutionary scenario driven by positive selection, compared to a 'random' mutational process (expected from neutral simulated data). Thus, mutation patterns denoting *homoplasy* and *stepwise evolution* are more likely to result from adaptation under selective scenarios, as transitioning amino acid changes towards novel apomorphic states can be considered rare events.

However, the number of sites scored under 'stepwise evolution' was higher (>25%) for both the simulated and expanded data. When visually verifying a subsample of these sites, a high proportion of false positives was detected (compared to a low number of false negatives). This is likely the result of intrinsic errors when adjusting our code to now exclude the MERS-CoV clade from the analyses. This observation further highlights the need to carefully custom-adjust code related to the input tree topology, and to the number of clades that are to be compared. Moreover, although our algorithm can be used as proxy for dataset reduction when searching for sites putatively denoting homoplasy and/or stepwise evolution, visual validation of the data is required to discard false positives and further validate positively scored mutational patterns (as shown in Supplementary Figure 3).

**Supplementary Figure 6. Position of a variable residue (genome site 23948) within the protein region preceding the S2’ cleavage site.**

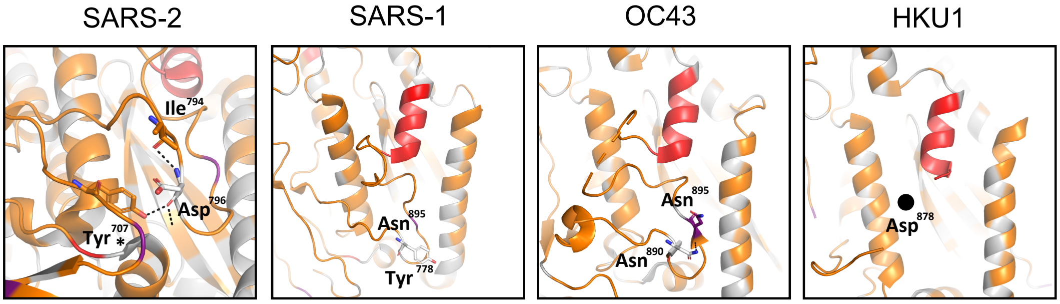


* *other protomer*

| | | |
|---|---|---|
| **SARS-2** | D796 | 793 PI K**D**FGGF**N**FSQILPDPS----------KPSKR\|SFIEDL 821 |
| **SARS-1** | Y778 | 775 TLK**Y**FGGF**N**FSQILPDPL----------KXTKR\|SFIEDL 803 |
| **OC43** | N890 | 887 VNF**N**VDDI**N**FXXVLGCLGSECSKASSR\|SAIEDL 919 |
| **HKU1** | D878 | 875 XHX**D**VDNIXFKSLXGCLGXXC--GSSSX\|SXXEDL 906 |

Schematic representation of the S protein structure for the four betacoronaviruses studied here. The cartoon panels show the protein structure preceding the S2’ cleavage site. The fusion peptide (indicated in red) is situated directly downstream of the S2’ cleavage site but is uncleaved in the structure. Residues of interest are shown in stick and ball representation. For SARS-CoV-2, genome position 23948 corresponds to a conserved D at residue 796 ($Asn^{796}$), whilst for SARS-CoV-1, this site corresponds to residue $Tyr^{796}$. At the homologous position, OC43 contains a conserved N (residue $Asn^{890}$), whilst HKU1 displays a conserved D (residue $Asp^{878}$). HKU1 residue $Asp^{878}$ is located within an unresolved region of the structure, and thus is represented using a black circle to indicate an approximate position. A conserved N-glycosylation sequon is shown in purple for three panels (corresponding to residue $N^{801}$ in SARS-CoV-2, $N^{783}$ in SARS-CoV-1 and $N^{895}$ in OC43). The asterisk for $Tyr^{707}$ in the S structure for SARS-CoV-2 indicates that this residue is located within a different protomer from the one containing the residues $D^{796}$ and $I^{794}$ leading to the fusion peptide. Below the panels, a consensus alignment is shown for the local amino acid sequence for each virus taxonomic group (see Methods Section 3). Amino acids of interest are highlighted in bold, whilst the conserved N-linked glycosylation sequon (NXT/S, in which X may be any amino acid except P) is underlined and the corresponding N is indicated in purple. For the HKU1 sequence, an ‘X’ within the sequence is either N or D, representing the loss of the N-glycosylation sequon within this virus taxonomic group. For the OC43 sequence, an ‘X’ within the N-glycosylation sequon can represent either an S or A, indicating either absence/ presence of the N-glycosylation sequon for different viral strains. The position of the S2’ cleavage site (R|S) is indicated preceding the fusion peptide (shown in red).

**Supplementary Figure 7. Distribution of positively (PSS) and negatively (NSS) selected sites within the SARS-CoV-2 Spike protein structure**

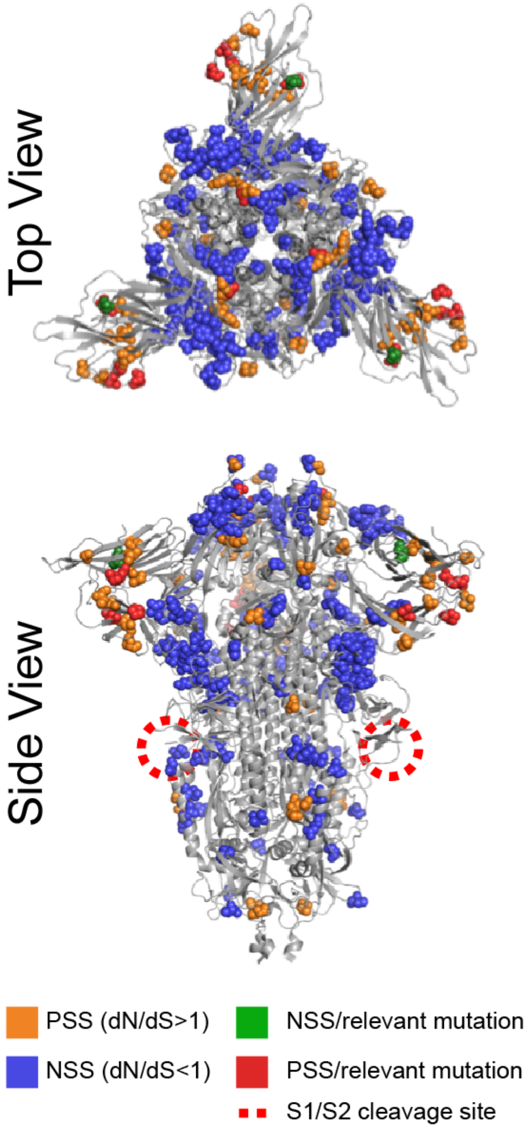


Top-down and side view of cartoon representation of the trimeric SARS-CoV-2 spike protein ectodomain (PDB: 6ZGI), showing the distribution of positively selected sites (PSS, scored under MEME), negatively selected sites (NSS, scored under FEL) (see Methods section 4, main text). PSS corresponding to the mutations of interest identified under our pipeline are shown in red, whilst the NSSs of interest are shown in green (PSS: residues 21, 25, 188 and 496, NSS: residue 122 in the SARS-CoV-2 spike protein numbering) (see Table 2). Other PSSs not scored under our pipeline are shown in orange, whilst other NSSs are shown in blue. Background residues are shown in grey. The S1/S2 cleavage site is indicated with a dashed circle in red. The furin cleavage site and surrounding sequence are not resolved within the structure (residues 677- 688; furin site RRAR are residues 682-685).

**Supplementary Table 2. PSS identified across all four virus species studied**

| | | PSS | |
|---|---|---|---|
| **Site †** | **Orf** | **MEME, % of branches dN/dS>1 *** | **p-value** |
| 293 | Orf1ab | 0.25 | 0.041 |
| 620 | Orf1ab | 0.16 | 0.002 |
| 638 | Orf1ab | 0.22 | 0.016 |
| 905 | Orf1ab | 0.20 | 0.042 |
| 2087 | Orf1ab | 0.24 | 0.027 |
| 2933 | Orf1ab | 0.30 | 0.036 |
| 3047 | Orf1ab | 0.02 | 0.004 |
| 3113 | Orf1ab | 0.16 | 0.000 |
| 3125 | Orf1ab | 0.03 | 0.005 |
| 3134 | Orf1ab | 0.05 | 0.050 |
| 3155 | Orf1ab | 0.63 | 0.025 |
| 3206 | Orf1ab | 0.10 | 0.015 |
| 3221 | Orf1ab | 0.14 | 0.043 |
| 3359 | Orf1ab | 0.37 | 0.049 |
| 3458 | Orf1ab | 0.03 | 0.035 |
| 4292 | Orf1ab | 0.21 | 0.004 |
| 4373 | Orf1ab | 0.23 | 0.025 |
| 5183 | Orf1ab | 0.17 | 0.032 |
| 5249 | Orf1ab | 0.17 | 0.014 |
| 5252 | Orf1ab | 0.44 | 0.048 |
| 5495 | Orf1ab | 0.11 | 0.013 |
| 5759 | Orf1ab | 0.21 | 0.002 |
| 5828 | Orf1ab | 0.03 | 0.050 |
| 6101 | Orf1ab | 0.02 | 0.031 |
| 6275 | Orf1ab | 0.30 | 0.018 |
| 6410 | Orf1ab | 0.23 | 0.027 |
| 6788 | Orf1ab | 0.16 | 0.048 |
| 6797 | Orf1ab | 0.06 | 0.041 |
| 8195 | Orf1ab | 0.12 | 0.045 |
| 9410 | Orf1ab | 0.24 | 0.010 |
| 9473 | Orf1ab | 0.02 | 0.003 |
| 10301 | Orf1ab | 0.21 | 0.024 |
| 10367 | Orf1ab | 0.10 | 0.034 |
| 11861 | Orf1ab | 0.04 | 0.026 |
| 12014 | Orf1ab | 0.03 | 0.005 |
| 13727 | Orf1ab | 0.02 | 0.003 |
| 17348 | Orf1ab | 0.06 | 0.029 |
| 17669 | Orf1ab | 0.13 | 0.022 |
| 17941 | Orf1ab | 0.05 | 0.030 |
| 18160 | Orf1ab | 0.24 | 0.031 |
| 18421 | Orf1ab | 0.13 | 0.037 |
| 18859 | Orf1ab | 0.03 | 0.044 |
| **19048** | **Orf1ab** | **0.03** | **0.036** |
| 19666 | Orf1ab | 0.13 | 0.010 |
| 21202 | Orf1ab | 0.03 | 0.034 |
| **21623** | **S** | **0.47** | **0.048** |
| **21635** | **S** | **0.57** | **0.048** |
| 21788 | S | 0.08 | 0.035 |
| 21803 | S | 0.10 | 0.032 |
| 21974 | S | 0.14 | 0.028 |
| 22037 | S | 0.99 | 0.036 |
| **22124** | **S** | **0.04** | **0.008** |
| 22127 | S | 0.02 | 0.028 |
| 22211 | S | 0.17 | 0.029 |
| 22331 | S | 0.72 | 0.020 |
| 22349 | S | 0.08 | 0.003 |
| 22823 | S | 0.03 | 0.022 |
| 22898 | S | 0.17 | 0.043 |
| 23045 | S | 0.19 | 0.005 |
| **23048** | **S** | **0.32** | **0.044** |
| 23129 | S | 0.07 | 0.040 |
| 23162 | S | 0.09 | 0.000 |
| 23615 | S | 0.02 | 0.035 |
| 23969 | S | 0.16 | 0.027 |
| 23972 | S | 0.07 | 0.032 |
| 24428 | S | 0.03 | 0.050 |
| 24908 | S | 0.15 | 0.043 |

**Supplementary Table 3. Location of other sites displaying evolutionary patterns putatively denoting homoplasy/stepwise evolution on protein regions of known function**

| SARS-CoV-2 reference genome coordinates * | ORF/ protein | Protein function | Structural Correspondence † ‡ | Structural proximity to known functional sites |
|---|---|---|---|---|
| **18121** | **Orf1ab/ nsp14** | **Exo RNAse** | **S28 in SARS-CoV-1 (PDB:5C8S)** | **Residue in the ExoN domain Proximal to the nsp10 interaction site., which cleaves terminal nucleotides during replication** |
| 20344 | Orf1ab/n sp15 | Endo RNAse | **H243** in SARS-CoV-2 (PDB:6WLC); **H242** in SARS-CoV-1 (PDB:2H85) | Within the NendoU catalytic domain, which cleaves non-terminal uracil nucleotides during replication |
| **21623** | **Spike ($S1^A$)** | **RBD** | **R21 in SARS-CoV-2; V25 in SARS-CoV-1; K29 in OC43; K28 in HKU1** | **Proximal to the $S1^A$ domain involved in receptor recognition for the embecoviruses** |
| **21635** | | | **P25 in SARS-CoV-2; N29 in SARS-CoV-1; P33 in OC43; V32 in HKU1** | **Proximal to the $S1^A$ domain involved in receptor recognition for the embecoviruses** |
| **23948** | Spike (S2) | Viral fusion | **D796 in SARS-CoV-2; Y778 in SARS-CoV-1; N890 in OC43; D878 in HKU1** | **Near the trimerization surface, which undergoes conformational rearrangements during viral fusion** |
| 24614 | | | **I1018** in SARS-CoV-2; **I1000** in SARS-CoV-1; **V1112** in OC43; **I1099** in HKU1 | In central helix domain, which undergoes conformational rearrangements during viral fusion |
| 24620 | | | **A1020** in SARS-CoV-2; **A1002** in SARS-CoV-1; **F1114** in OC43; **A1101** in HKU1 | In central helix domain, which undergoes conformational rearrangements during viral fusion |
| 24632 | | | **L1024** in SARS-CoV-2; **L1006** in SARS-CoV-1; **Q1118** in OC43; **R1105** in HKU1 | In central helix domain, which undergoes conformational rearrangements during viral fusion |

* For SARS_CoV_2|Wuhan-Hu-1|MN908947 reference sequence. Sites in bold refer to those referred to in Table 1 and main text.
† Structures of the relevant protein (domains) have not been solved for all four betacoronaviruses studied here.
‡ Available structures used in this study: PDB 2W2G, PDB 5C8S, PDB 6WLC, PDB 5I08, PDB 6OHW, PDB 6ACC and PDB 6VXX (see Methods section 5)
